# Splicing modulators impair DNA damage response and induce killing of cohesin-mutant MDS/AML

**DOI:** 10.1101/2022.09.26.509430

**Authors:** Emily C Wheeler, Benjamin J E Martin, William C Doyle, Rebecca A Gorelov, Melanie Donahue, Johann C Jann, Omar Abdel-Wahab, Justin Taylor, Michael Seiler, Silvia Buonamici, Roger Belizaire, Karen Adelman, Zuzana Tothova

**Affiliations:** Department of Medical Oncology, Dana-Farber Cancer Institute; Boston, MA 02215, USA; Broad Institute of MIT and Harvard University; Cambridge, MA 02142, USA, Cambridge, MA 02142, USA; Department of Biological Chemistry and Molecular Pharmacology, Blavatnik Institute, Harvard Medical School; Boston, MA 02115, USA; Ludwig Center at Harvard, Harvard Medical School; Boston MA 02115, USA; Human Oncology and Pathogenesis Program, Memorial Sloan Kettering Cancer Center; New York, NY 10021, USA; Division of Hematology, Department of Medicine, Sylvester Comprehensive Cancer Center at the University of Miami Miller School of Medicine; Miami, FL 33136, USA; H3 Biomedicine, Inc.; 300 Technology Square, Cambridge, MA 02139, USA; Department of Pathology, Dana-Farber Cancer Institute; Boston, MA 02215, USA

## Abstract

Splicing modulation is a promising treatment strategy pursued to date only in splicing-factor mutant cancers; however, its therapeutic potential is poorly understood outside of this context. Like splicing factors, genes encoding components of the cohesin complex are frequently mutated in cancer, including myelodysplastic syndromes (MDS) and secondary acute myeloid leukemia (AML), where they are associated with poor outcomes. Here, we show that cohesin mutations are biomarkers of sensitivity to drugs targeting the splicing-factor SF3B1 (H3B-8800 and E-7107). We identify drug-induced alterations in splicing and corresponding reduced gene expression of a large number of DNA repair genes, including *BRCA1* and *BRCA2,* as the mechanism underlying this sensitivity in cell line models, primary patient samples and patient-derived xenograft (PDX) models of AML. We find that DNA damage repair genes are particularly sensitive to exon skipping induced by SF3B1 modulators given their long length and large number of exons per transcript. Furthermore, we demonstrate that treatment of cohesin-mutant cells with SF3B1 modulators not only results in impaired DNA damage response and accumulation of DNA damage, but it significantly sensitizes cells to subsequent killing by PARP inhibitors and chemotherapy, and leads to improved overall survival of PDX models of cohesin-mutant AML *in vivo*. Our findings expand the potential therapeutic benefits of SF3B1 splicing modulators to include cohesin-mutant MDS and AML, and we propose this as a broader strategy for therapeutic targeting of other DNA damage-repair deficient cancers.

**One Sentence Summary:** We identify an unexpected effect of SF3B1 splicing inhibitors on regulation of DNA damage repair genes and show efficacy of combination treatment in cohesin-mutant MDS and AML.

## INTRODUCTION

Cohesin is a multi-subunit protein complex that is essential for sister chromatid cohesion, three dimensional chromosome organization and gene regulation (*1*). It forms a ring around DNA, with three structural subunits, SMC1A, SMC3, and RAD21 bound to either STAG1 or STAG2 proteins. Cohesin is one of the most frequently mutated protein complexes in cancer and mutations in genes encoding components of the cohesin ring and its modulators are frequent and recurrent drivers in myeloid malignancies. These include myelodysplastic syndromes (MDS) and acute myeloid leukemia (AML) where cohesin mutations are associated with poor overall survival (*2–5*). Loss-of-function mutations in *STAG2* are the most common mutations found in cohesin subunits and result in altered cellular functions including longer loop extrusion, altered gene expression, replication fork stress, and accumulation of DNA damage (*6–10*).There are currently no targeted therapies available for patients with cohesin-mutant cancers. However, our recent work describing DNA damage accumulation and the genetic dependency on DNA repair proteins in cohesin-mutant MDS and AML (*6*) has led to a Phase 1B clinical trial of single agent PARP inhibitor talazoparib treatment (ClinicalTrials.gov identifier NCT03974217).

*SF3B1* is the most frequently mutated gene in MDS, and splicing factor mutations in general comprise the most common class of genetic alterations found in MDS and secondary AML (sAML) (*11–14*). Widespread alternative splicing and reliance on proper splicing function is a common feature of cancer cells that has been implicated in disease progression and exploited for therapeutic targeting (*15–18*). Cancer cells accumulate mis-spliced RNAs through a variety of mechanisms including the acquisition of hotspot point mutations or alterations in the expression levels of splicing factor proteins (*19–21*). Therapeutic targeting of the spliceosome is of particular interest in myeloid malignancies where splicing factor mutations are common, such as MDS, AML and chronic myelomonocytic leukemia (CMML) (*22*). H3B-8800 and E-7107 are first-in-class compounds that modulate splicing function through targeting of the most commonly mutated splicing factor SF3B1, which is a component of the U2snRNP involved in branch point recognition (*23*). These compounds have shown promise in preclinical studies and phase 1/2 clinical trials where treatment with H3B-8800 resulted in transfusion independence in a subset of MDS patients with splicing factor mutations (*24–26*). However, the mechanisms mediating response to SF3B1-modulators are not well understood and it remains unclear whether these drugs will provide therapeutic benefit beyond splicing-factor mutant cancers.

In this work, we identify widespread mis-splicing of DNA repair genes in cells treated with SF3B1-modulators, leading to an accumulation of DNA damage. Further, treatment of cohesin-mutant cancer cells with H3B-8800 or E-7107 results in an increased therapeutic vulnerability to chemotherapeutic agents targeting DNA repair. We use human-cell derived *in vitro* and *in vivo* models of cohesin-mutant AML and patient samples from splicing factor-mutant patients enrolled in the H3B-8800 clinical trial to demonstrate that mis-splicing of DNA damage repair genes results in reduced expression of DNA damage response proteins and accumulation of DNA damage in cells. While the splicing changes caused by SF3B1-modulators are genotype agnostic, we find that cohesin-mutant cells are specifically targeted for killing due to their dependence on proper DNA damage repair for survival (*6, 27*). We leverage this vulnerability to develop a new therapeutic strategy in which treatment of cohesin-mutant cancer cells with low-dose splicing modulation dramatically increases the sensitivity to compounds that induce DNA damage or inhibit proper DNA damage repair, including PARP inhibitors and a variety of chemotherapeutic agents. Thus, we propose that splicing modulation in combination with inhibition of DNA damage repair may be an effective therapeutic strategy to target cohesin-mutant MDS/AML, and other cancers that exhibit an accumulation of DNA damage or selective reliance on the DNA repair machinery.

## RESULTS

### Cohesin-mutant cells are sensitive to SF3B1-targeting compounds

To generate cell line models of cohesin-mutant AML, we used CRISPR-Cas9 editing to establish a panel of isogenic single cell-derived clones from parental U937 and K562 cells which are wild type for all cohesin complex subunits and modulators (*6*). We engineered these cells to contain loss-of function mutations in *STAG2* and *SMC3* commonly found in patients (Figure 1A). We had previously used these cell lines to perform genome-wide CRISPR screens with the goal of identifying mutant-specific genetic dependencies and potential therapeutic vulnerabilities (*6*). In addition to a preferential dependency of *STAG2*-mutant cells on STAG1 and DNA damage and replication machinery, we identified *STAG2*-mutant cells to be more sensitive to knockout of genes involved in mRNA-processing. To further explore the interaction of these mutations in primary patients, we analyzed co-mutation patterns in patients in our clinical cohort of MDS patients as well as publicly available datasets. We identified that mutations in *STAG2* tend not to co-occur with *SF3B1* mutations, suggestive of synthetic lethality (Figure 1B, S1A), (*11, 12*). To further probe this possibility, we tested if loss-of-function cohesin mutations are tolerated in the background of the most frequently mutated *SF3B1* codon K700E. We performed CRISPR-mediated depletion of *STAG2* or *SMC3* in isogenic *SF3B1-K700E* and *SF3B1*-wild type (*SF3B1*-*K700K*) K562 cells and found that loss of any of the cohesin subunits reduced cell viability of *SF3B1-K700E* mutant relative to wild type cells (Figure S1B). These data were consistent with our observation that *STAG2* and *SF3B1* mutations co-occur less frequently than predicted by chance and raised the possibility that cohesin-mutant cells may be vulnerable to splicing modulation.

**Fig. 1.**
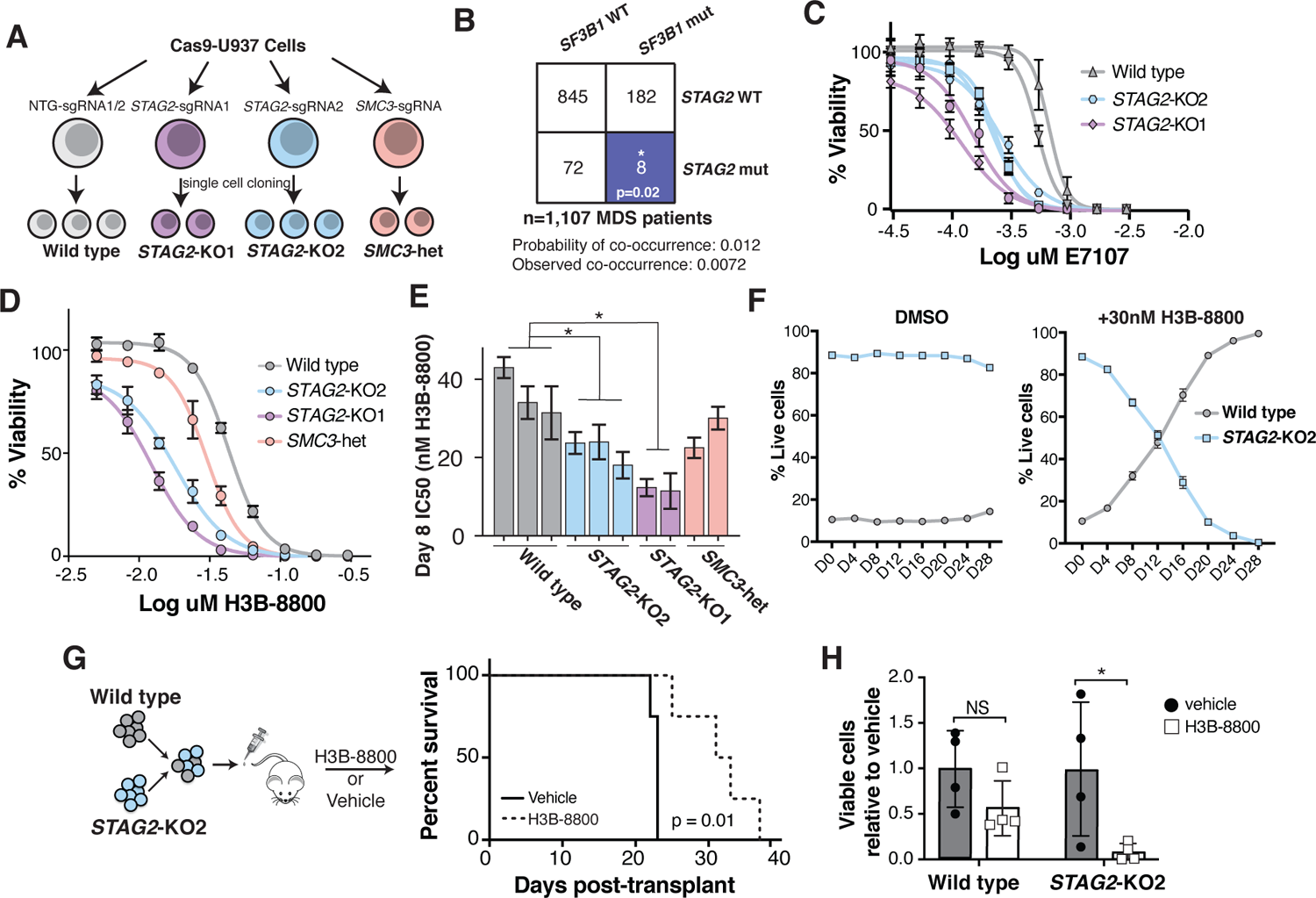
Cohesin-mutant cells are sensitive to SF3B1-targeting compounds. (**A**) Schematic of isogenic AML cell lines used in this study. U937 cells expressing Cas9 were nucleofected with single-guide RNAs (sgRNAs) targeting *STAG2, SMC3* or non-targeting (NTG) sgRNAs. Two independent sgRNAs were used for *STAG2* (KO1 and KO2) and NTG, and a single sgRNA was used to target SMC3. Independent single cell-derived clones were used as biological replicates in this study. (**B**) Co-occurrence of *SF3B1* and *STAG2* mutations in a cohort of MDS patients from the Dana-Farber Cancer Institute. Expected and observed probably of co-occurrence are listed. Blue color indicates a significant mutual exclusive relationship between *SF3B1* and *STAG2* mutations. *p<0.05 (Z-test). WT = wild type, mut = mutant. (**C**) Drug dose-response curves of E-7107–treated WT and *STAG2*-U937 clones on Day 12 of treatment. Error bars represent SD of measurements of three technical replicates. (**D**) Drug dose response curves of a representative set of wild type and cohesin-mutant U937 cells treated with H3B-8800 for 8 days. Error bars represent SD of measurements in technical triplicates. (**E**) Quantification of IC_50_ among biological replicates (n=2 or 3) of wild type and cohesin-mutant U937 cells on day 8 of treatment with H3B-8800 tested in technical triplicates. *STAG2*-KO1 and *STAG2*-KO2 clones have a significantly lower IC_50_ than wild type cells with H3B-8800 treatment (Kruskal-Wallis with post-hoc test, p=0.05). Error bars represent the 95% confidence interval of the IC_50_ calculated from technical triplicates of each cell line. (**F**) Competition assay with wild type (mCherry) and *STAG2*-KO2 (GFP) U937 cells mixed in a 1:10 ratio in the presence of DMSO or H3B-8800 (30nM) *in vitro*. % Live GFP+ or mCherry+ cells were determined using flow cytometry. Error bars represent SD of measurements of technical triplicates. (**G**) Schematic of the *in vivo* drug treatment of NSGS mice injected with wild type (mCherry+) and *STAG2*-KO2 (GFP+) U937 cells mixed in a 1:1 ratio. Treatment with H3B-8800 or vehicle control was started 7 days post-transplant. Kaplan-Meier survival analysis was performed; p=0.01. n=4 mice per group. (**H**) Leukemia burden in mice treated with H3B-8800 or vehicle was assessed in the spleen of animals at the time of sacrifice. % Live GFP+ or mCherry+ cells were determined using flow cytometry. Mean ± SD is shown. P<0.05 (Student’s t-test). n=4 mice per group.

To understand if splice-modulating compounds may provide therapeutic benefit in cohesin-mutant MDS and AML, we treated cohesin-mutant U937 and K562 AML cell lines with the SF3B1-targeting compounds H3B-8800 and E-7107 at increasing concentrations *in vitro*. We found that cohesin-mutant cells were more sensitive to both splicing modulators as compared to the isogenic, wild type controls (Figures 1C-E, Figures S1C-E). In competition assays, *STAG2*-mutant cells were outcompeted by wild type cells in the presence of low dose H3B-8800 even when present at a 10-fold excess (Figure 1F). We also performed *in vivo* competition experiments using mouse xenograft models injected with a mixture of wild type and *STAG2*-mutant AML cells and observed both selective killing of the *STAG2*-mutant cells by H3B-8800 and a survival benefit in animals treated with the drug (Figures 1G-H). These results demonstrate that cohesin-mutant AML cell lines exhibit sensitivity to SF3B1-targeted splicing modulation and led us to further interrogate the mechanism driving this response.

To determine if the sensitivity to SF3B1-modulating compounds in cohesin-mutant cells is driven by common splicing alterations in cohesin-mutant and *SF3B1 K700E*-mutant cells at baseline, we compared the splicing events altered upon mutation in each of these models. First, we compared RNA-seq from U937 cell clones of each cohesin-mutant genotype to wild type control cells (Figure 1A). We quantified a total of 3,740 splicing events that are significantly altered in any of the three cohesin-mutant lines. K-means clustering of these events revealed highly similar patterns of splicing among the cohesin-mutant cells, with many exon skipping and exon inclusion events in common (Figure S1F). Next, we analyzed publicly available RNA-Seq data from K562 cells with *SF3B1 K700E* mutations (*26*) or *STAG2*-KO (*28*) and identified 2,874 splicing events that are significantly altered in either genotype as compared to wild-type K562 cells. K-means clustering performed on these events demonstrated little overlap between the splicing changes observed in the *SF3B1 K700E*-mutant versus *STAG2*-KO cells (Figure S1G). Based on the shared splicing changes identified across the cohesin-mutant U937 cell lines, and clear differences in splicing observed in cohesin-mutant versus *SF3B1 K700E*-mutant K562 cells, we suggest that the reliance of cohesin-mutant cells on proper splicing does not reflect a common set of disrupted splicing events in cohesin-mutant and *SF3B1 K700E*-mutant cells. Instead, we propose that disruption of splicing with SF3B1 inhibitors targets cancer cells through a shared therapeutic vulnerability that exists downstream of splicing regulation in both cohesin-mutant and *SF3B1 K700E*-mutant cells.

### H3B-8800 treatment induces mis-splicing and downregulation of DNA damage repair genes

To understand the mechanism by which splicing modulating drugs target cohesin-mutant cells, we treated our panel of ten isogenic U937 cell lines with increasing concentrations of the SF3B1-modulator H3B-8800 and quantified splicing and gene expression changes relative to DMSO treatment using total RNA-Seq. We hypothesized that treatment with H3B-8800 may target cohesin-mutant cells by either (i) selectively modulating splicing of essential genes in cohesin-mutant but not wild type cells, or (ii) leading to similar splicing changes in all genotypes but targeting genes that are selective dependencies for the survival of cohesin-mutant cells. We identified over 13,000 splicing changes in H3B-8800 treated cells relative to DMSO, the majority of which were exon skipping events that are alternatively spliced in a dose-dependent manner (Figures 2A, Figure S2A-B). We used k-means clustering to group splicing events based on the percent change in splicing and observed that H3B-8800-induced splicing changes are largely conserved in all cells, including both wild type and cohesin-mutant (Figure 2B). In general, splicing changes induced by H3B-8800 occur in exons that are normally fully included (PSI = 1) and are spliced out uniquely in the presence of the drug. Conversely, we find that introns retained upon H3B-8800 treatment are usually fully spliced out (PSI = 0) (Figure 2C). Thus, rather than reflecting alternative splicing, the changes observed in H3B-8800-treated cells often represent mis-splicing and the formation of aberrant mRNA products. In agreement with this model, the median gene expression of H3B-8800 target genes decreased in a dose-dependent manner suggesting that drug-induced alterations in splicing result in impaired gene expression (Figure S2C). These results support that H3B-8800 causes similar splicing changes in all genotypes, and that the preferential killing of cohesin-mutant cells by H3B-8800 is driven by mis-splicing of genes that are selectively essential for survival in cohesin-mutant, but not wild type cells.

**Fig. 2.**
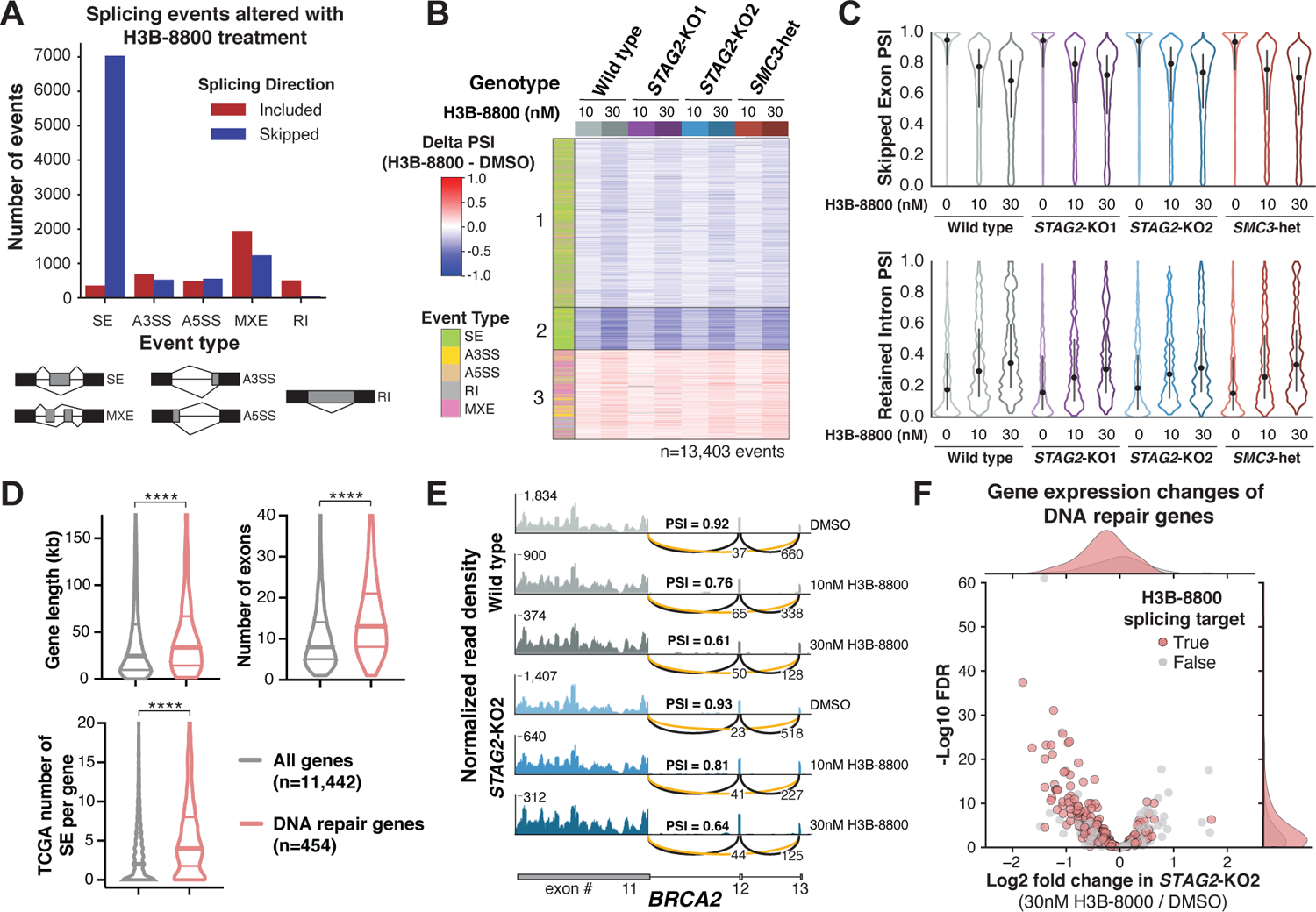
H3B-8800 treatment induces mis-splicing and downregulation of DNA damage repair genes. (**A**) Total number and directionality of splicing alterations induced by 6-hr H3B-8800 treatment in all U937 cell lines. Events are categorized by event type and direction of regulation in H3B-8800–treated relative to DMSO-treated cells. (SE = skipped exon, A3SS = alternative 3’ splice site, A5SS = alternative 5’ splice site, MXE = mutually exclusive exon, RI = retained intron). (**B**) Heatmap of delta PSI scores for H3B-8800–regulated exons (from panel A) across all conditions separated into 3 k-means clusters. Each comparison consists of two (*STAG2*-KO1, *SMC3*-heterozygous) or three (*STAG2*-KO2, wild type) independent single cell-clones for each concentration of drug compared to DMSO controls within the same genotype. Columns are organized by genotype and concentration of H3B-8800. Color bar on the left indicates the type of splicing event that was called. PSI (percent spliced in). (**C**) Violin plots showing the distribution of PSI scores for H3B-8800–regulated skipped exons (top) and retained introns (bottom) under each treatment condition tested. Dot represents the median, and bars extend from the first to third quartile range. PSI (percent spliced in). (**D**) Violin plots depicting the gene length, number of exons, and number of skipped exons in TCGA comparing DNA repair genes (n=454) to all expressed protein-coding genes (n=11,442). Horizontal lines in violin plots depict the median and 1^st^ and third quartiles. **** p < 0.0001, Mann-Whitney test. (**E**) RNA-Seq normalized read density and splice junction track of exon skipping in *BRCA2* exon12 from one representative replicate of wild type and *STAG2*-KO2 cells treated with DMSO, 10nM, or 30nM H3B-8800 for 6 hours. Average PSI scores from 3 biological replicate samples of exon12 are shown. Average number of reads supporting exon skipping (orange line) and exon inclusion (black line) are reported. PSI (percent spliced in). (**F**) Volcano plot of gene expression changes in DNA repair genes in *STAG2*-KO2 cells treated with 30nM H3B-8800 relative to DMSO-treated controls. Average log2 fold change of 3 biological replicates of *STAG2*-KO2 versus wild type U937 cells is shown. DNA repair genes that contain H3B-8800–regulated splicing changes are highlighted in red.

Having established that H3B-8800–induced splicing changes are shared by cohesin wild type and mutant cells, we next investigated candidate genes whose mis-splicing could lead to preferential killing of cohesin-mutant cells. We started by quantifying enrichment of gene ontology (GO) terms among the genes in each cluster of H3B-8800-induced splicing changes from Figure 2B. DNA repair was the only GO category among the top five enriched terms in all three clusters (Figure S2D, Table S1). As discussed above, cohesin-mutant cells are known to accumulate DNA damage and rely on proper DNA repair for survival (*6, 27*), making this category of genes particularly interesting for further study.

Notably, DNA damage repair genes tend to be significantly longer, contain more exons per transcript, and are more often found to have skipped exons compared to all other expressed genes in the Cancer Genome Atlas (TCGA) data set (Figures 2D, S2E-F) (*29*). We thus propose that these features make DNA damage repair genes highly dependent on accurate splicing for their expression, rendering them more sensitive to splicing changes induced by H3B-8800. An example of H3B-8800–induced exon skipping is shown for exon12 of *BRCA2* which is skipped in 40-45% of transcripts in wild type and *STAG2*-KO2 cells treated with 30nM H3B-8800 for 6 hours (Figure 2E). Among the 454 annotated DNA damage repair genes (*30*) expressed in U937 cells, 70% contain H3B-8800–induced splicing changes and ∼40% of those contain a reduction in gene expression in a dose-dependent manner (Figures 2F, S2G-H). Therefore, the splicing changes observed in DNA damage repair genes in cells treated with H3B-8800 may lead to loss of proper DNA repair, a process that is shared between wild type and mutant cells, but particularly critical for survival of cohesin-mutant cells.

### Mis-splicing of DNA repair genes alters protein function and results in accumulation of DNA damage

Among the top mis-spliced and downregulated genes with H3B-8800 treatment were *BRCA1* and *BRCA2* (Figures 3A, S3A), proteins that play a critical role in DNA repair and whose loss of function is known to correlate with sensitivity to Poly (ADP-ribose) polymerase (PARP) inhibitors. We confirmed that at the protein level, cells treated with H3B-8800 nearly completely lost expression of both BRCA1 and BRCA2 72 hours after treatment (Figure 3B). Mis-splicing in other DNA damage repair genes, such as *CHEK2,* did not lead to major changes in gene expression, but resulted in loss of normal protein function by splicing out amino acid residues in key functional domains (Figures 3C, S3A). H3B-8800–induced skipping of *CHEK2* exon2 splices out the T68 amino acid targeted for phosphorylation and initiation of downstream signaling, as well as exon10, a portion of the kinase domain required for phosphorylation of its substrates (Figure 3C).

**Fig. 3.**
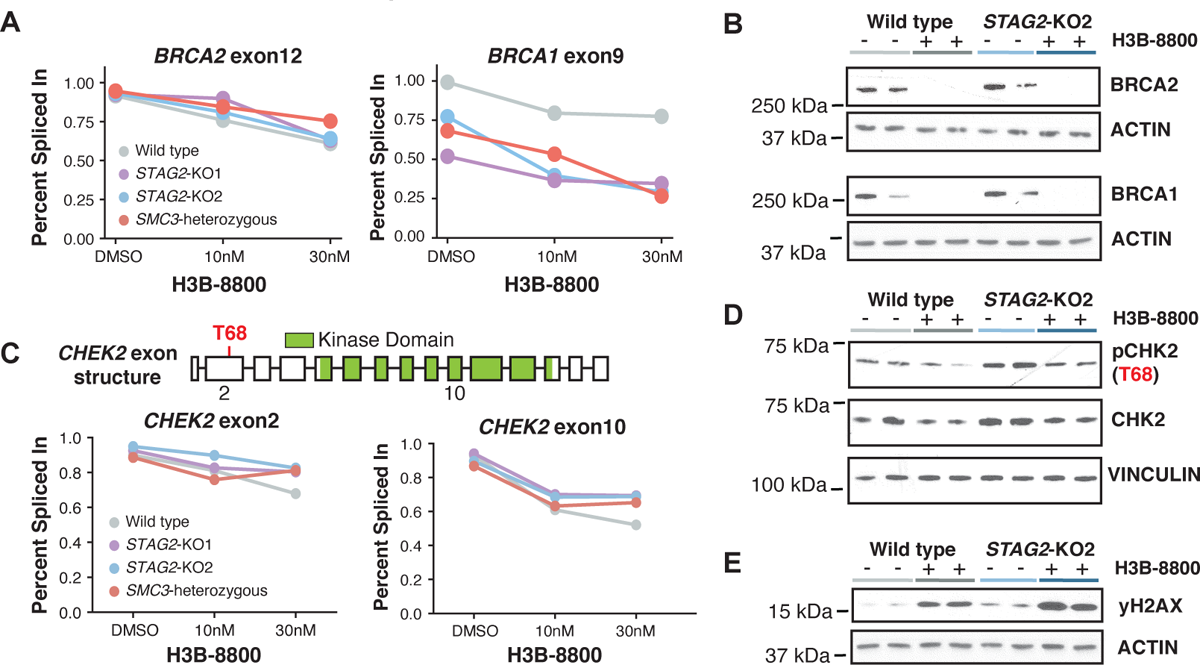
Mis-splicing of DNA repair genes alters protein function and results in accumulation of DNA damage. (**A**) Average PSI scores of *BRCA2* exon12 and *BRCA1* exon9 are plotted for each genotype and drug condition. Datapoints represent the mean of biological triplicates (wild type and *STAG2*-KO2) or duplicates (*STAG2*-KO1 and *SMC3*-heterozygous) for each treatment condition. (**B**) Western blot analysis of BCRA1 and BRCA2 protein levels in wild type and *STAG2*-KO2 cells treated with 50nM H3B-8800 or DMSO for 3 days. Each lane represents an independent single-cell clone of U937 cells transduced with a non-targeting sgRNA (wild type), or sgRNA targeting *STAG2*. Actin was used as a loading control. (**C**) Schematic of *CHEK2* exon structure (top) with the Thr68 phosphorylation residue highlighted in red and the annotated kinase domain shown in green. Percent Spliced In (PSI) scores of exon2 and exon10 are shown for each treatment condition. Datapoints represent the mean of biological triplicates (wild type and *STAG2*-KO2) or duplicates (*STAG2*-KO1 and *SMC3*-heterozygous) for each treatment condition. (**D**) Western blot analysis of pCHK2 and total CHK2 protein levels in wild type and *STAG2*-KO2 cells treated for 3 days with 50nM H3B-8800 or DMSO. Each lane represents an independent single-cell clone of U937 cells transduced with a non-targeting sgRNA (wild type), or sgRNA targeting *STAG2*. Vinculin was used as a loading control. (**E**) Western blot analysis of γH2AX protein levels in wild type and *STAG2*-KO2 cells treated for 3 days with 50nM H3B-8800 or DMSO. Each lane represents an independent single-cell clone of U937 cells transduced with a non-targeting sgRNA (wild type), or sgRNA targeting *STAG2*. Actin was used as a loading control.

At the protein level, H3B-8800–treatment both depletes total CHK2 levels and pCHK2 abundance, thus impairing the ability to initiate proper DNA repair through signaling of the CHK2 DNA damage checkpoint (Figure 3D). As evidence of accumulating DNA damage following H3B-8800 treatment, western blotting for the marker of DNA damage γH2AX showed substantial increase in signal in both cohesin-mutant and wild type U937 cells (Figure 3E). These results support a model in which H3B-8800 inhibits DNA repair through mis-splicing of key DNA repair genes in a genotype-independent manner, resulting in the accumulation of DNA damage.

### Splicing modulation sensitizes cohesin-mutant AML cell lines to killing by talazoparib and chemotherapy

One prediction of our model is that treatment of cells with H3B-8800 or E-7107 impairs DNA damage repair and therefore renders cells more sensitive to subsequent treatment with DNA damage repair inhibitors or DNA damage-inducing agents. Having established significant drug-induced mis-splicing and loss of expression of BRCA1 and BRCA2, two key regulators of homology-directed repair (HDR) and biomarkers of PARP inhibitor sensitivity, we first tested whether sequential treatment of cells with an SF3B1 modulator followed by the PARP inhibitor talazoparib potentiates killing of cohesin-mutant cells. Pre-treatment of cohesin wild type U937 cells with either of the two available SF3B1 modulators, H3B-8800 or E-7107, followed by treatment with talazoparib, resulted in a minimal decrease in cell viability (Figures 4A-B). Pre-treatment of *STAG2*-KO U937 cells with DMSO, followed by treatment with talazoparib, recapitulated increased sensitivity of *STAG2*-KO versus wild type cells to talazoparib as we have previously reported (*6*). However, pre-treatment of *STAG2*-KO U937 cells with either of the two splicing modulators significantly increased their sensitivity to talazoparib with up to a 10-fold increase in killing (Figures 4A-B, S4A). Furthermore, cells accumulated more DNA damage when subjected to sequential treatment of H3B-8800 followed by talazoparib than with either drug alone, as assessed by γH2AX western blot (Figures S4B-C).

**Fig. 4.**
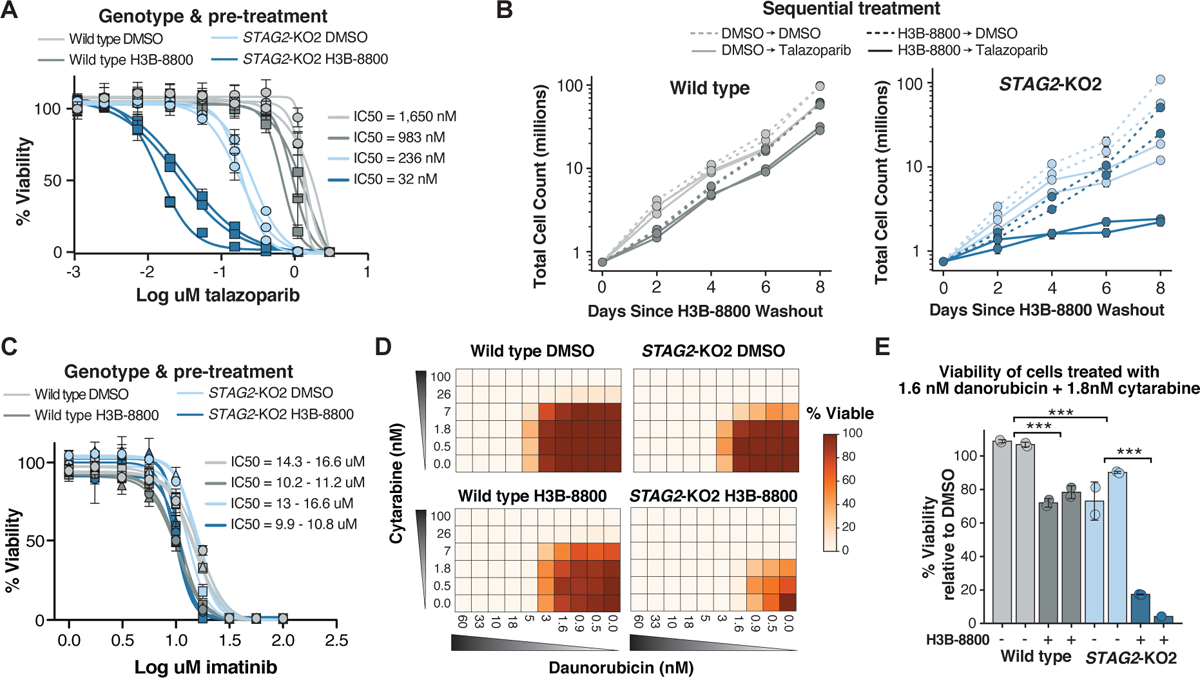
Splicing modulation sensitizes cohesin-mutant AML cell lines to killing by talazoparib and chemotherapy. (**A**) Drug dose-response curves of wild type and *STAG2*-KO2 cells pre-treated with 50nM H3B-8800 or DMSO for 3 days, followed by drug washout and 8 days of treatment with talazoparib. Error bars represent SD of technical triplicate measurements for each biological triplicate sample (n=3 per genotype and condition). (**B**) Growth curves depicting total number of wild type or *STAG2*-KO2 cells pre-treated with DMSO or H3B-8800 (50nM) for 3 days, followed by treatment with talazoparib (50nM) or DMSO for 8 days. Error bars represent SD of technical duplicate measurements for each biological replicate sample (n=2 per genotype and condition). (**C**) Drug dose-response curves of wild type and *STAG2*-KO2 cells pre-treated with 50nM H3B-8800 or DMSO for 3 days, followed by drug washout and 8 days of treatment with imatinib. Error bars represent SD of technical triplicate measurements for each biological replicate sample (n=2 per genotype and condition). (**D**) Heatmap of cell viabilities for wild type and *STAG2*-KO2 cells pre-treated with DMSO (top) or 50nM H3B-8800 (bottom) for 3 days, followed by treatment with a combination of daunorubicin and cytarabine for 8 days. Cell viabilities are normalized to DMSO-treated controls on each plate (0nM cytarabine and 0nM daunorubicin). Values shown are the average of two technical replicate samples for one representative biological replicate sample. (**E**) Bar plot of percent viable cells following combination treatment with 1.6nM daunorubicin and 1.8nM cytarabine relative to DMSO-treated controls. Cells received either 3 days of 50nM H3B-8800 (dark bars) or DMSO control (light bars) prior to chemotherapy. Data points show technical replicates n=2 per condition (except biological replicate 2 of H3B-8800 treated *STAG2*-KO2 cells n=1). Each bar represents an independent biological replicate sample. ***p<0.0001, student’s T-test.

We next wanted to determine if treatment of cells with an SF3B1 splicing modulator could sensitize cells to additional DNA damage-inducing agents, such as chemotherapy. We tested a panel of chemotherapeutic drugs, including single agent and combination treatment of cytarabine and daunorubicin commonly used in the treatment of AML, as well as etoposide, cisplatin and mitomycin C. Pre-treatment of cells with H3B-8800 followed by treatment with any of these chemotherapeutic agents resulted in increased killing of both cohesin-mutant and wild type cells, with preferential sensitivity of cohesin-mutant cells over wild type cells (Figure S4D). In contrast, pre-treatment with H3B-8800 had no effect on viability of cells treated with imatinib, an ABL kinase inhibitor used to treat chronic myeloid leukemia (CML) and CML blast crisis, which does not induce DNA damage (Figure 4C). Furthermore, treatment of cells with combination of daunorubicin and cytarabine, the standard of care regimen for AML, resulted in 80-90% depletion of cohesin-mutant cells after a single dose pre-treatment with H3B-8800 compared to only 25-30% killing in wild-type cells (Figures 4D-E, S4E). Therefore, short-term treatment with a splicing modulator may be an effective method to increase the therapeutic window of many DNA damage-inducing chemotherapeutic agents, particularly in subsets of cancer with an underlying DNA damage repair defect, such as cohesin-mutant AML.

### Low dose splicing modulation combined with talazoparib or chemotherapy targets PDX AML *in vivo*

To test if treatment with splicing modulators sensitizes primary patient samples to DNA damage repair inhibitors *in vivo*, we generated three serially transplantable patient-derived xenograft (PDX) models of AML from two different human *STAG2*-mutant and one cohesin-wild type AML samples (Figures 5A, S5A) (*6*). We first tested if treatment with E-7107 could reproduce the effect on splicing and downregulation of DNA damage repair genes in *vivo*. To do this, we performed RNA-Seq on human bone marrow cells isolated from one of the *STAG2*-mutant and one cohesin-wild type AML PDX model following 3 or 5 days of E-7107 treatment. Consistent with our cell line results, we observed widespread exon skipping and splicing changes that are conserved at the event-level among genes that were detected in the PDX cells (Figures 5B-C, S5B-C). DNA repair genes, including BRCA1 and BRCA2 were downregulated in both PDX models tested (Figures 5D, S5D-G). These data support that our initial observations from AML cell lines are conserved in primary patient cell-derived xenograft models.

**Fig. 5.**
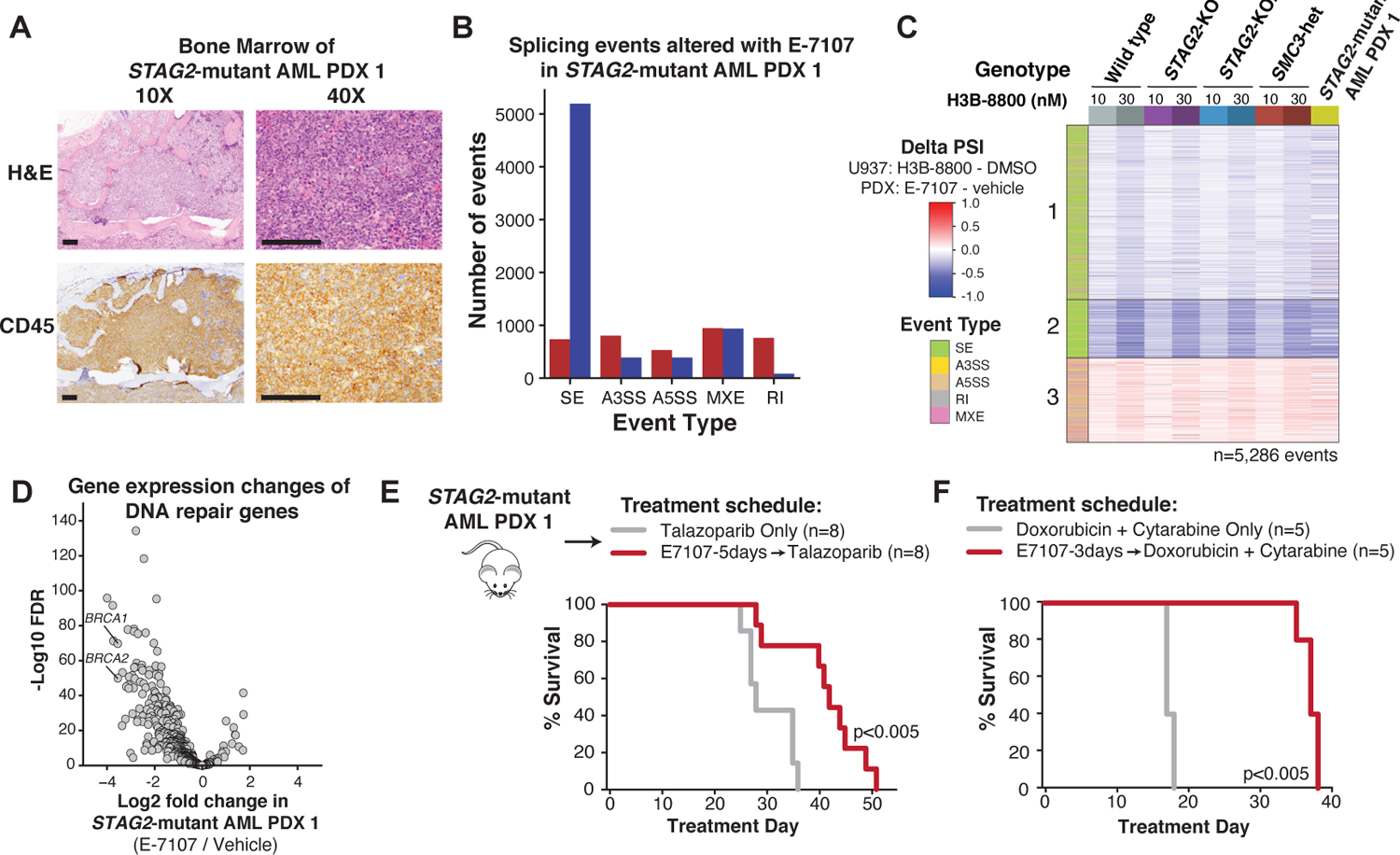
Low-dose splicing modulation combined with talazoparib or chemotherapy targets PDX AML *in vivo.* (**A**) Morphologic evaluation of bone marrow of *STAG2*-mutant AML1 patient derived xenograft shows infiltration with human leukemia blasts. Images were stained using H&E and hCD45-targeting antibody and imaged at 10x and 40x (scale bar: 0.125mm) original magnification. (**B**) Total number and directionality of significant splicing alterations differentially called in *STAG2*-mutant human AML1 PDX cells isolated from bone marrow of NSGS mice treated with E-7107 compared to vehicle for 5 days *in vivo.* Splicing events are categorized by event type and direction of regulation in E-7107 versus vehicle treated mice. (SE = skipped exon, A3SS = alternative 3’ splice site, A5SS = alternative 5’ splice site, MXE = mutually exclusive exon, RI = retained intron). N=3 mice per condition. (**C**) Heatmap of delta PSI scores for H3B-8800–regulated exons called from U937 cells (Figure 2B) that are expressed in *STAG2*-mutant AML PDX1 treated with E-7107 *in vivo*. Each comparison consists of two (*STAG2*-KO1, *SMC3*-heterozygous) or three (*STAG2*-KO2, wild type, *STAG2*-mutant PDX) independent biological replicates compared either DMSO or vehicle treated controls. Color bar on the left indicates the type of splicing event that was called, column colors are labeled by genotype and drug treatment on the right. (**D**) Volcano plot depicting differential gene expression of DNA repair genes in *STAG2*-mutant human AML1 PDX cells isolated from the bone marrow of NSGS mice treated with E-7107 versus vehicle for 5 days *in vivo*. N=3 mice per condition. (**E**) Schematic of *in vivo* E-7107 drug treatment and survival analysis of *STAG2*-mutant AML1 PDX model (other mutations include *BCOR/RUNX1/U2AF1/DNMT3A*). Treatment of mice assigned to two treatment arms was initiated 3 weeks after bone marrow transplantation: talazoparib only (n=8) or E-7107 for 5 days followed by talazoparib (n=8). Survival data was combined from two independent experiments. P<0.005 (log-rank test). (**F**) Survival analysis of *STAG2*-mutant AML1 PDX mice treated with 3 days of E-7107 followed by combination chemotherapy (doxorubicin + cytarabine), or combination chemotherapy alone (n=5 mice per arm). P<0.005 (log-rank test).

Having established that splicing targets and downregulation of DNA repair genes with E-7107 treatment was conserved *in vivo*, we next treated *STAG2*-mutant AML PDX models sequentially with splicing modulation or chemotherapy to determine if the addition of splicing modulation can improve overall survival. First, we treated a cohort of mice with 3 or 5 days of E-7107 followed by talazoparib, compared to mice treated with talazoparib alone. We observed a significant benefit in overall survival among both E-7107-containing treatment arms, with the longer splicing modulator pre-treatment time contributing to the strongest benefit in overall survival (Figures 5E, S5H). Pre-treatment with E-7107 also delayed the disease-associated thrombocytopenia (Figure S5I). We treated a second *STAG2*-mutant AML PDX model with 5 days of E-7107 followed by talazoparib, or talazoparib alone. Notably, this model was treated at a more advanced disease state with 20-50% human circulating cells. The E-7107 treatment arm reduced the human cells in the blood by 20 – 40% after 5 days of treatment and contributed to a significant benefit in overall survival (Figures S5K-L). We next investigated the effect of SF3B1 splicing inhibition followed by combination chemotherapy. We observed significantly improved overall survival of *STAG2*-mutant AML PDX mice after treatment with 3 days of E-7107 followed by combination chemotherapy of doxorubicin and cytarabine, reminiscent of standard induction treatment of AML in humans (Figures 5F, S5J). These results confirm that low-dose SF3B1 splicing modulation provides a therapeutic benefit in combination with PARP inhibition or chemotherapy *in vivo* and should be considered as a potential treatment strategy for cohesin-mutant cancers.

### Splicing changes and downregulation of DNA repair genes are conserved in MDS/AML patients

Having established that SF3B1-modulator treatment leads to mis-splicing of DNA damage repair genes and sensitization of cohesin-mutant cells and PDX models to DNA damage-inducing drugs, we next wanted to confirm that these effects were conserved in primary human samples. We first interrogated splicing and gene expression changes among the eleven genes that were assessed using a custom nanostring panel as on-target biomarkers in the Phase 1 clinical trial of H3B-8800 in splicing factor-mutant MDS and AML (Steensma et al., 2021). Ten out of eleven genes were expressed in U937 cells and eight contained H3B-8800–regulated splicing changes that met our stringent threshold for significance (Figure S6A). Moreover, seven of the eight mis-spliced genes showed a dose-dependent effect on gene expression (Figures S6B). The remaining two genes, *FBXW5* and *DYNLT1*, contained multiple introns that are retained in a dose-dependent manner throughout the gene body, but at levels that failed to meet our thresholds (Figure S6C-D). These data confirm on-target activity of H3B-8800 in our *in vitro* AML cell line models.

Next, we wished to address whether H3B-8800 treatment in patients led to mis-splicing of DNA damage repair genes. Since comprehensive RNA-Seq analysis on patients from the H3B-8800 clinical trial had not been reported, we performed total RNA-sequencing on peripheral blood samples collected from patients pre-treatment and 2-4 hours after receiving their first oral dose of H3B-8800 (Figure 6A, Table S2). Splicing changes relative to the pre-treatment sample were quantified for three patients independently, given the variable doses administered. Consistent with our *in vitro* data, exon-skipping was the most common alternative splicing event detected following H3B-8800 treatment, and the number of events increased in a dose-dependent manner among the three patients (Figure 6B). We then asked if the drug-induced splicing changes identified in U937 cells (as shown in Figure 2B) were conserved in patient samples. Strikingly, we observed a dose-dependent increase in splicing alterations that mirrors the splicing changes observed in U937 cells (Figures 6C, S6E). In the patient who received the highest dose of H3B-8800, 35% of expressed DNA repair genes contained at least one drug-induced splicing change, including the same event that was detected in *BRCA1* in U937 cells (Figure 6D). To examine whether splicing changes that occurred within 2-4 hours of H3B-8800 administration could already impact levels of gene expression, we performed a paired differential expression analysis comparing each patient pre- and post-treatment. As expected from the short time frame of treatment, changes in gene expression were generally small in magnitude. However, a trend towards downregulation of DNA repair genes was observed in all patients following treatment with H3B-8800 (Figure 6E). These results confirm that there is conservation at the splicing event-level with H3B-8800 treatment *in vivo* and *in* vitro and demonstrate that DNA repair genes are targeted in patient samples.

**Fig. 6.**
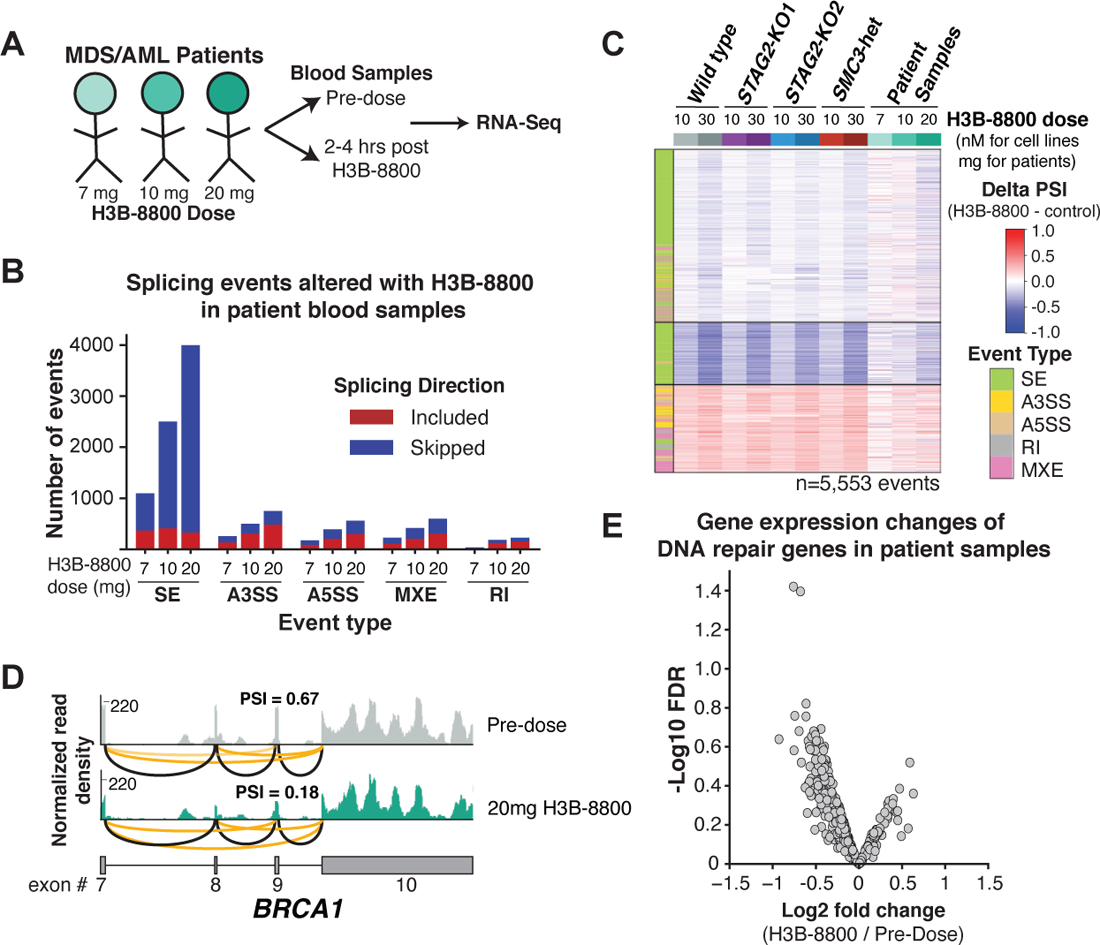
Splicing changes and downregulation of DNA repair genes are conserved in MDS/AML patients. (**A**) Schematic of samples collected from MDS/AML patients treated with 3 different doses of H3B-8800 on clinical trial (ClinicalTrials.gov identifier NCT02841540). (**B**) Total number and directionality of significant splicing alterations differentially called in each patient sample pre- and post-H3B-8800. Patients are sorted on the x axis according to increasing doses of H3B-8800. Splicing events are categorized by event type and direction of regulation in H3B-8800 versus pre-treatment sample. (SE = skipped exon, A3SS = alternative 3’ splice site, A5SS = alternative 5’ splice site, MXE = mutually exclusive exon, RI = retained intron). (**C**) Heatmap of delta PSI scores for H3B-8800–regulated splicing changes called from U937 cells (Figure 3A) that are expressed in patient samples. Patient samples are sorted by increasing dose of H3B-8800 received. Color bar on the left indicates the type of splicing event that was called, column colors are labeled by genotype and drug treatment. (**D**) RNA-Seq normalized read density and splice junction track of exon skipping in *BRCA1* exon 9 from the pre- and post-treatment sample in the patient who received 20mg of H3B-8800. Black lines indicate constitutive splicing junctions, orange lines indicate splice junctions that contain exon skipping. PSI=percent spliced in. (**E**) Volcano plot depicting differential gene expression of DNA repair genes from a paired analysis of all patients pre- and post-H3B-8800 treatment.

## DISCUSSION

Our studies establish a role for RNA splicing modulation in the therapeutic targeting of cohesin-mutant MDS and AML. We demonstrate increased sensitivity of cohesin-mutant cells to splice-modulating drugs that target the SF3B complex and propose to include SF3B1 modulation in combination with chemotherapy or PARP inhibition to target cohesin-mutant cells. While E-7107 and H3B-8800, the first two SF3B1 modulators, were first designed to treat MDS and AML patients harboring splicing factor mutations, this work suggests that MDS and AML patients with cohesin mutations may also derive therapeutic benefit from this class of drugs.

We characterize a mechanism by which H3B-8800 and E-7107 induce cell death through mis-splicing of DNA repair proteins, a class of genes that are enriched for long genes with many exons. We show that treatment with H3B-8800 results in accumulation of γH2AX and reduced function and expression of key DNA damage response genes including CHK2, BRCA1, and BRCA2, which we have previously described as an important genetic dependency in cohesin-mutant cells (*6*). While there are currently no targeted therapies approved for cohesin-mutant patients, clinical testing of single-agent PARP inhibition in cohesin-mutant MDS and AML is ongoing (ClinicalTrials.gov identifier NCT03974217). We propose to expand this effort by using a dual-agent strategy in which low-dose splicing modulation is used to further sensitize cohesin-mutant cells to PARP inhibition or standard chemotherapy. Importantly, we show that this approach increases survival in primary patient-derived *STAG2*-mutant AML xenografts.

While splicing modulators that target the SF3B complex were first developed to treat patients with splicing factor mutations, current research suggests that they may have widespread utility beyond this subset of cancer patients. Other cancer types, including *MYC*-driven triple negative breast cancer, T-ALL with upregulation of SRSF7, CLL irrespective of *SF3B1* mutation status, and aggressive prostate cancer, have shown efficacy of SF3B1-targeting drugs in preclinical models (*31–34*). Our findings suggest that cohesin-mutant cancers may similarly benefit from splice-modulating therapies, particularly those targeting the SF3B complex. It remains to be seen if this sensitivity will extend to other types of splice-modulating drugs that are undergoing clinical development, including SRPK inhibitors, RBM39 degraders, and PRMT inhibitors (*22, 35*).

It is important to note that none of the patient samples we analyzed from the H3B-8800 clinical trial contained mutations in *STAG2* or other cohesin proteins. In addition, the changes in splicing and downregulation of DNA repair genes are strongly conserved among both cohesin wild-type and mutant PDX models and cell line models, with the same splicing changes present irrespective of the genotype. These findings suggest that a general treatment strategy of splicing modulation followed by PARP inhibition or chemotherapy is likely to be relevant in a wide range of tumors that rely on proper DNA damage repair for survival (*36*). These include *BRCA*-mutant breast and ovarian cancers characterized by a “BRCA-ness” phenotype of dysfunctional HDR and resulting sensitivity to PARP inhibition (*37, 38*).

Recent work has suggested that spliceosome-mutant MDS cell line models are also sensitive to PARP inhibition, making this class of cancers another candidate for dual splicing and PARP inhibition therapy (*39*). A major challenge with single agent PARP inhibitor treatment is acquired resistance through escape mechanisms in which cells repair HDR defects (*38*). However, targeting cancer dependency pathways with multiple, non-overlapping agents is a proven strategy to prevent acquisition of resistance. We propose low dose splicing modulation followed by PARP inhibition or standard chemotherapy as a new therapeutic strategy to be considered in cohesin-mutant MDS and AML and suggest a potential benefit of this strategy in other cancer types that are deficient in proper DNA repair. In summary, our study identifies a critical connection between cohesin mutations and splicing modulation in AML, creating a novel therapeutic strategy for these patients with very limited treatment options and poor outcomes.

## MATERIALS AND METHODS

### Experimental Design

The objective of this study was to determine the role of RNA splicing modulation in the survival of cohesin-mutant MDS/AML. Isogenic cell line models derived as multiple independent single-cell clones per genotype were used for all *in vitro* work. 2-3 independent biological replicates were used for each individual experiment, all of which were performed in technical replicates or triplicates. For *in vivo* studies, the primary endpoint was overall survival, and the secondary endpoint was depletion of human cohesin-mutant cells. Animals were sacrificed according to pre-determined endpoints of 20% weight loss from the beginning of treatment, or a body condition score of 2 or lower. For animal studies, treatment groups were assigned to normalize blood counts per group prior to treatment. The studies were not performed in a blind fashion.

### U937 and K562 cell lines and culture

U937 and K562 cell lines used in this study were previously published (*6*). Cells were grown in RMPI + 10% FBS + 1% Pen/Strep/Glutamine at 37°C with 5% CO_2_. Mycoplasma testing is routinely performed on cells in culture.

### U937 cell transplantation mouse model

8-10 week old female NSGS mice (NOD-SCID; IL2Rγ null; Tg(IL3, CSF2, KITL)) were obtained from The Jackson laboratory, strain 013062). Recipient mice were sublethally irradiated using a Gamma cell irradiator (Best Theratronics) at a dose of 250 Rads and injected by tail vein with 500,000 GFP+ or mCherry+ STAG2-mutant or WT U937 cells. All mice were housed in a pathogen-free animal facility in microisolator cages and experiments were conducted according to an IACUC approved protocol at the Dana-Farber Cancer Institute.

### PDX mouse models

The PDX models used in this study were generated using adult female NSGS mice (NOD-SCID; IL2Rγ null; Tg(IL3, CSF2, KITL), The Jackson Laboratory, strain 013062), which were irradiated using an gamma cell irradiator (Best Theratronics) with 200 rads ≤24 hours prior to transplant. The *STAG2*-mutant AML PDX1 sample was from a male patient obtained with informed consent under Dana-Farber/Harvard Cancer Center Institutional Review Board (IRB)-approved protocol #19152. Unsorted mononuclear cells were collected from the bone marrow aspirate. Dana-Farber Cancer Institute’s Rapid Heme Panel Sequencing analysis revealed the following mutation and variant allele frequencies: STAG2 ENST00000371160.1:c.2096+1G>A 66.8%, BCOR ENST00000397354.3:c.4138C>T 63.3%, BCORL1 ENST00000218147.7:c.3583A>T 48.7%, RUNX1 ENST00000300305.3:c.805+1G>T 31.6%, U2AF1 ENST00000291552.4:c.101C>T 30.7%, DNMT3A ENST00000321117.5:c.2597+1delG 29.9%, ATM ENST00000278616.4:c.8269G>A 4.0%. Unsorted mononuclear cells were thawed, resuspended in Hanks’ Balanced Salt Solution (Thermo Fisher Scientific 14025076), and injected via tail vein into two 31-week old NSGS mice (“P0”). For the *STAG2*-mutant AML PDX1, P0 mice were sacrificed when they became morbid, and engraftment of human cells in the bone marrow and spleen was measured with flow cytometry (CD45-PE-Cy7 antibody, BD Biosciences). Single-cell suspensions were prepared from the bone marrow, and spleen, the cells were washed, red blood cells lysed prior to injection into another cohort of 10-week old NSGS mice using the protocol described above (“P1”). Mice from this cohort were monitored daily and sacrificed when they became morbid. Cells were collected as described above and frozen in 90% FBS, 10% DMSO freezing media. Cells from P1 mice were thawed in PBS + 1%FBS, washed and resuspended in Hanks’ Balanced Salt Solution for injection of 1.3 – 2 million cells per mouse via tail vein into 8-week old NSGS mice (“P2”). Dana-Farber Cancer Institute’s Rapid Heme Panel Sequencing analysis was performed on the bone marrow cells from P2 mice to confirm the mutation status of the engrafted clone as follows: BCOR c.4240C>T (p.Q1414*) 100% VAF, BCORL1 c.3583A>T (p.K1195*) 81.5% VAF, DNMT3Ac.2597+1delG (splice site) 37.7% VAF, RUNX1 c.805+1G>T (splice site) 20.3% VAF, STAG2c.2096+1G>A (splice site) 97.9% VAF, U2AF1 c.101C>T (p.S34F) 32.9% VAF. Mice from the P2 generation were used for all experiments done in this study.

The cohesin wild type PDX model was generated from an AML patient with informed consent under Dana-Farber/Harvard Cancer Center Institutional Review Board (IRB)-approved protocol #01-026. Unsorted mononuclear cells were collected from a leukapheresis sample were frozen and sequencing analysis from Dana-Farber Cancer Institute’s Rapid Heme Panel Sequencing analysis revealed the following mutation and variant allele frequencies: DNMT3A c.2645G>A (p.R882H) 49.6% VAF, NPM1 c.859_860insTCTG (p.W288Cfs*12) 43.9% VAF, PTPN11 c.1504T>G (p.S502A) 1.8% VAF, FLT3-ITD. Cells were thawed in PBS + 1%FBS, washed and resuspended in Hanks’ Balanced Salt Solution for injection of 1.3 – 2 million cells per mouse via tail vein injection for all mice used in this study.

### CRISPR lentiviral transduction

sgRNAs targeting *STAG2*, *SMC1A*, *SMC3*, or non-targeting (NTG), were cloned into lentiCRISPRv2 (Addgene #52961) as described (http://genome-engineering.org/gecko/wp-content/uploads/2013/12/lentiCRISPRv2-and-lentiGuide-oligo-cloning-protocol.pdf). Lentivirus was produced in 293TL cells, and cells were transduced by spin infection at 2200rpm for 90 minutes at 37C in the presence of 4ug/mL polybrene. Following completion of antibiotic selection, cells were seeded for downstream assays as described.

### *In vivo* H3B-8800 dosing of U937-transplant model

Mice were dosed daily with 8mg/kg H3B-8800 or an equivalent volume of vehicle in 0.5% methylcellulose solution daily by oral gavage, starting on Day 7 post-transplantation; treatment continued for 15-30 days until the mice were sacrificed upon disease development. 4 recipients were injected per experimental group. Mice were weighed daily, and the daily dose was not administered if the body weight dropped by ≥10% from the previous day. Animals were monitored daily for evidence of disease and were sacrificed when morbid. Following sacrifice, mice were examined for presence of *STAG2*-mutant cells in the bone marrow and spleen with flow cytometry. Single-cell suspensions were made from bone marrow, and spleen, washed, red blood cells were lysed, and samples were frozen in 90% FBS, 10% dimethylsulfoxide (DMSO) until analysis.

### Flow cytometry for U937 cells

U937 cells were single cell sorted with a FACSAriaII instrument (Becton Dickinson, Mountain View, CA) after DAPI staining for viability (Thermo Fisher Scientific). mCherry+ and GFP+ U937 cells used for in vivo transplant studies and in vitro competition assays were sorted with a MoFlo Astrios EQ sorter (Beckman Coulter) or Sony SH800S Sony Cell Sorter. Readout of *in vitro* competition assays was performed using CytoFLEX (Beckman Coulter).

### U937 cells treated with H3B-8800 for RNA-Seq

U937 cells were treated with H3B-8800 at 10nM, 30nM or DMSO for 6 hours. 1 million cells were collected in 350ul of Trizol and flash frozen. RNA was extracted with Zymogen Direct-Zol columns. 100ng of input RNA was used for library preparation with the KAPA RNA hyper prep kit with RiboErase treatment according to manufacturer’s instructions. Libraries were sequenced on the Illumina NOVASeq in PE100 mode to a depth of 100M reads per sample at the Dana-Farber Cancer Institute.

### *In vitro* drug treatment and *in vitro* competition assays

All drug dose response assays were conducted in 96 well plates. For U937 and K562 cell lines, 10,000 cells were plated per well at 0.05×10^6^ cells/ml. Cells were split 1:4 by volume every 4 days and redosed with fresh medium supplemented with fresh drug every 4 days. For pre-treatment with DMSO, H3B-8800 (50nM), or E-7107 (1.14nM), cells were cultured in bulk for 3 days before plating in 96-well format for secondary drug dosing. Viability was measured every 4 days using the CellTiter-Glo luminescent cell viability assay (Promega G7573). A D300e drug dispenser (Tecan) was used for drug dosing in all experiments performed in 96 well at the concentrations indicated for each drug. Drug dose-response curves were fitted and IC50 concentrations were calculated using GraphPad Prism. All drugs used in this study are listed in the key resources table.

For cell counting assays at single drug dose concentrations, cells were cultured in 6-well dishes at the recommended concentration by cell type and split to equal concentration of cells for passaging. Drugs were redosed at the time of passage (3 or 4 days). Viable cell counts were measured in duplicate with a ViCell XR cell viability analyzer (Beckman Coulter).

For competition experiments, mCherry-labeled wild type and GFP-labeled STAG2-KO2 U937 cells were mixed in a 1:10 ratio and plated in 96-well plates with 3 technical replicates. Cells were passaged and split 1:4 by volume every 4 days and redosed with fresh medium supplemented with fresh H3B-8800 every 4 days. A fraction of the cells were stained for viability with DAPI, and % mCherry+DAPI- and GFP+DAPI-cells were determined by flow cytometry.

### E-7107 *in vivo* drug treatment and RNA extraction for RNA-Seq

Three weeks after transplant, *STAG2-*mutant AML PDX1 mice were bled, and a complete blood count was performed with an Element HT5 analyzer (Heska). Mice were assigned to treatment groups (n=3 per group) based on platelet counts and treatment began 4 weeks post-injection for 5 days. For the cohesin wild type AML PDX, mice were bled 5 weeks after transplant and %hCD45 cells were measured in the circulation by flow cytometry. Mice were assigned to treatment groups (n=2 per group) based on %hCD45+ cells in the peripheral blood and treatment began 8 weeks post-injection for 3 days. E-7107 (4mg/kg in 10% Ethanol, 5% Tween80 in sterile saline solution) or an equivalent volume of Vehicle, was delivered daily via tail vein injection. Four hours following the final injection, mice were sacrificed and single-cell suspensions were made from bone marrow, cells were washed and red blood cells were lysed. Samples were stained and sorted for viable, human CD45+ cells (BD 557748) directly into RLT lysis buffer supplemented with b-Mercaptoethanol on a BD FACSAria II cell sorter. RNA extraction was performed using a Qiagen RNeasy Mini Kit. 50ng of RNA was used as input for library preparation with the KAPA mRNA HyperPrep kit. Libraries were sequenced on the Illumina NOVASeq in PE100 mode to a depth of 100M reads per sample at the Dana-Farber Cancer Institute.

### E-7107 + talazoparib *in vivo* drug treatment

For the *STAG2*-mutant AML PDX1, mice were bled three weeks after transplant and a complete blood count was performed with an Element HT5 analyzer (Heska). Mice were assigned to treatment groups based on platelet counts and treatment began 4 weeks post-injection with a 5-days-on, 2-days-off treatment schedule. For the *STAG2-*mutant AML PDX2, mice were bled monthly to monitor %hCD45 cells in circulation by flow cytometry and to perform a complete blood count with an Element HT5 analyzer (Heska). 6 months after transplant, mice were assigned to treatment groups based on %hCD45 cells in the peripheral blood. For the first 5 days of treatment, mice received both a tail vein injection of E-7107 (4mg/kg in 10% Ethanol, 5% Tween80 in sterile saline solution) or Vehicle, and oral gavage of Talazoparib (0.25mg/kg in 0.5% methylcellulose), or methylcellulose alone depending on the treatment group. The second week of treatment continued only with Talazoparib dosing until the time of sacrifice. Mice were weighed daily, and drug was withheld if the body weight dropped by ≥15% from the starting mouse weight. Mice were monitored daily and were sacrificed when morbid or when they had lost 20% of the starting body weight. Engraftment of human cells in the bone marrow, spleen, and peripheral blood was determined with flow cytometry.

### E-7107 + doxorubicin and cytarabine *in vivo* drug treatment

*STAG2-*mutant AML PDX1 mice were bled three weeks after transplant and a complete blood count was performed with an Element HT5 analyzer (Heska). Mice were assigned to treatment groups based on platelet counts and treatment began 4 weeks post-injection. Treatment days 1 – 3 mice received either E-7107 (IV, 4mg/kg in 10% Ethanol, 5% Tween80 in sterile saline solution) or vehicle only control. On days 4 – 8, mice in the chemotherapy arm received 5 days of cytarabine (IP 10mg/kg in sterile saline solution), and 3 days of doxorubicin (IV, 1mg/kg in PBS) given in combination with cytarabine on days 4-6. The E-7107 arm received 5+3 days of cytarabine and doxorubicin on days 6-10 following 2 days of rest after E-7107. Mice were weighed daily and sacrificed when morbid or when they had lost 20% of the starting body weight. Engraftment of human cells in the bone marrow, spleen, and peripheral blood was determined with flow cytometry.

### Flow cytometry for hCD45 detection in blood samples from PDX models

Peripheral blood samples were collected with retro-orbital bleeding in EDTA coated tubes. Red blood cells were lysed twice with Lysing Buffer (BD Biosciences 555899) and remaining cells were stained for human CD45 (BD 557748). Percent hCD45+ cells in the blood was determined with flow cytometry.

### Western Blot

Cells were washed in cold 1X PBS (Corning CellGro) and lysed for 15-30min on ice using NP-40 lysis buffer (150mM NaCl, 50mM Tris pH 7.5 (Thermo Fisher Scientific 15567027), 1% NP40 (Roche 11332473001), 5% glycerol (Sigma-Aldrich G5516), and 100X Halt Protease and Phosphatase Inhibitor Cocktail (Thermo Fisher Scientific 78447) in water). Lysates were quantified using Pierce BCA Protein Assay Kit (Thermo Fisher Scientific 23225) and reduced with 6X Laemmli SDS-Sample Buffer (Boston BioProducts BP-111R) and 10X NuPAGE Sample Reducing Agent (Thermo Fisher Scientific NP0009) for 5min at 95°C. Samples were run on NuPAGE 4-12% Bis-Tris Protein Gels (Invitrogen NP0321 and NP0316) in NuPAGE MOPS SDS Running Buffer 20X (Life Technologies NP0001) or NuPAGE MES SDS Running Buffer 20X (Life Technologies NP0002) and transferred onto Nitrocellulose Blotting Membranes (0.45μm, Invitrogen LC2001) using 10X Tris/Glycine Buffer (Bio-Rad 1610734) and 20% Methanol (VWR TXBJ349664LBRI) in water. Gels were transferred at 30V overnight at 4°C. Membranes were blocked with 5% nonfat dry milk (Bio-Rad 1706404XTU) in Tris Buffered Saline with Tween-20 (Cell Signaling Technology, 9997) for 30-60min at room temperature. Incubation with primary antibody was performed overnight at 4°C in 5% milk in TBST. Primary antibodies and concentrations include: Vinculin (V9131 sigma, 1:1000), CHK2 (MP 05-649, 1:5000), pCHK2 (CST2661, 1:1,000), y-H2AX (CST9718, 1:1000), Actin (CST8H10D10, 1:10,000), BRCA2 (OP95, 1:500), BRCA1 (MS110, 1:1000). Membranes were washed 3 times for 15min each in TBST and incubated in Anti-mouse IgG, or Anti-rabbit IgG secondary as in 5% milk in TBST for 2hrs at room temperature. Membranes were washed an additional 3 times for 15min each in TBST and visualized after a 5min incubation with SuperSignal West Dura Extended Duration Substrate (Thermo Scientific, 34075), or SuperSignal West Pico Chemiluminescent Substrate (Thermo Fisher Scientific, 34078).

### RNA-Seq analysis

U937 RNA-seq samples were aligned to the hg19 genome, and PDX RNA-seq seq samples were aligned to a combined hg19 and mm10 genome using STAR (version 2.7.3a) (*40*). PDX reads uniquely aligning to hg19 were used to quantify gene expression and splicing. Gene counts were generated using featurecounts function of the Rsubread package (version 2.0.1) (*41*) aligning reads to the Ensembl (GRCh37.87) basic transcript annotations. Differentially expressed genes were calculated using DESeq2 (version 1.26.0) (*42*), normalizing samples by sequencing depth. Genes with a log_2_ foldchange in expression greater than 0.5 and an adjusted p-value less than 0.001 were called as differentially expressed.

For U937 expressed gene annotations, Ensembl annotations (GRCh37.87) were filtered for basic annotations. Transcript expression in each RNA-seq sample was quantified using kallisto (version 0.45.1) (*43*). Transcripts with expression greater than 0.5 TPM in at least two samples were retained, giving 37,389 transcripts corresponding to 20,120 genes. These expressed transcript annotations were then used for analyzing differentially expressed genes and alternative splicing. We used our get gene annotations pipeline (*44*) to call the dominant transcript isoforms for each gene, using the RNA-seq data as input. Briefly, this pipeline clusters annotated TSSs within 100bp and summed TPM values in each cluster. The cluster with the highest expression was selected as the dominant cluster, and the transcript with the highest expression was selected as the dominant TSS. Then for transcripts stemming from the dominant TSS cluster, we cluster transcript end sites (TESs), and, similar to TSSs, selected the dominant TES cluster and TES. The transcript with the highest TPM was then selected as the dominant transcript model for counting the number of exons per gene. We focused our analysis on protein coding genes, and this yielded 11,442 dominant transcript models for further analysis.

### Splicing analysis

Alternative splicing was quantified using rMATS (version 4.1.0) (*45*) considering junction-spanning reads only. All comparisons to generate delta PSI scores are as listed in the manuscript using biological replicates for each condition. For *STAG2*-mutant AML PDX1 samples with *in vivo* E-7107 treatment, bam files from three replicate samples of the E-7107 treated arm were merged to obtain one replicate to increase coverage. The E-7107 treated arm had dramatic depletion of the human cells, and therefore many fewer reads were obtained in those samples, so merging was performed to increase read depth on splice junctions. The vehicle treated arm had sufficient coverage in each replicate and therefore were kept separate for analysis.

We used custom scripts available on github to process the raw output files and remove overlapping splicing calls on the same event, selecting only the event call with the most inclusion reads (https://github.com/YeoLab/rbp-maps/blob/master/preprocessing_scripts/subset_rmats_junctioncountonly.py). Next, unique events were further filtered to only include those with a minimum of 10 reads supporting both the inclusion and skipped isoforms in at least one condition. Significantly regulated splicing events (absolute delta PSI > 5%, FDR < 0.05) were then called for each comparison of interest.

For clustering analysis, events that were called significant in any comparison were considered and deltaPSI was quantified from those events across all comparisons that are included in the clustering analysis. kMeans clustering was performed with the with an optimal k and the deltaPSI scores were plotted per cluster in a heatmap. For the analysis comparing all U937 cells treated with 10nM and 30nM H3B-8800, k=4 was used for the clustering and one cluster was dropped from the final analysis for a total of 3 displayed clusters. The removed cluster contained sporadic splicing events that were only significant in one cell type or drug concentration and were not representative of the on-target effects of H3B-8800 in cells.

### Patient Sample RNA-Seq

Blood samples were collected from patients immediately before and 2-4 hours following their first dose of H3B-8800. RNA was extracted from whole blood, quantified and 100ng was used as the input for total RNA-Seq. Reads were aligned to hg19 as described above. Differential gene expression was calculated with DESeq2 using a paired analysis, comparing the pre- and post-H3B-8800 sample per patient with the design formula of ∼patient + treatment. Alternative splicing was quantified with rMATS for each patient individually comparing the pre- and post-H3B-8800 treatment sample. All filtering steps to call significantly regulated splicing changes were performed as described above.

### Statistical Analysis

Statistical analysis was performed in GraphPad PRISM8, and with the Scipy Stats module in Python (v1.5.0). The specifics of statistical tests used and samples size for each experiment are described in the figure legends. No outliers were removed from data analysis.

## List of Supplementary Materials

Fig. S1. Cohesin-mutant cells are sensitive to SF3B1-targeting compounds.

Fig. S2. H3B-8800 treatment induces mis-splicing and downregulation of DNA damage repair genes.

Fig. S3. Mis-splicing of DNA repair genes alters protein function and results in accumulation of DNA damage

Fig. S4. Splicing modulation sensitizes cohesin-mutant AML cell lines to killing by talazoparib and chemotherapy

Fig. S5. Low-dose splicing modulation combined with talazoparib targets PDX AML *in vivo*

Fig. S6. Splicing changes and downregulation of DNA repair genes are conserved in MDS/AML patients

Table S1. Gene Ontology of H3B-8800 Target Genes

Table S2. Patient Sample Information

## Acknowledgments

The authors would like to thank the members of the Tothova and Adelman labs for helpful discussion and feedback on this work. We thank Suzan Lazo, Michael Buonopane, and James Marcucella at the Dana-Farber Flow Cytometry core for help with cell sorting and analysis, Maura Berkeley and Zach Hebert from the Molecular Biology Core Facility at Dana-Farber for preparation of RNA-Seq libraries and sequencing. We thank Elodie Hatchi and Anna Idelevich from the Livingston lab for providing BRCA1 and BRCA2 antibodies and helpful discussion on this work.

The work of Z.T. and J.T. was supported by the The Doris Duke Charitable Foundation. The work of Z.T. was supported by The Burroughs Welcome Fund and The Edward P. Evans Foundation. The work of Z. T. and K.A. was supported by the Ludwig Center at Harvard and the Quadrangle Fund for Advancing and Seeding Translational Research (QFSATR) at Harvard Medical School. ECW is supported by the NIH Ruth L. Kirschstein F32 postdoctoral award (F32HL159905). BJM is supported by a CIHR Banting Postdoctoral Fellowship and QFASTR.

## Funding

Doris Duke Charitable Foundation (ZT, JT) Burroughs Welcome Fund (ZT)

Edward P. Evans Foundation (ZT) Ludwig Center at Harvard (ZT, KA)

Quadrangle Fund for Advancing and Seeding Translational Research (QFASTR) at Harvard Medical School (ZT, KA, BJM)

NIH Ruth L. Kirschstein F32 postdoctoral award F32HL159905 (ECW) CHIR Bandint Postdoctoral Fellowship (BJM)

## Author contributions

Conceptualization: ECW, BJM, KA, ZT Methodology: ECW, BJM, KA, ZT

Investigation: ECW, BJM, WCD, RAG, MD, JCJ, MS, RB

Visualization: ECW, BJM, WCD Funding acquisition: KA, ZT Resources: OAW, SB Supervision: OAW, SB, KA, ZT Writing – original draft: ECW

Writing – review & editing: BJM, KA, ZT

## Competing interests

MS and SB were employees of H3 Biomedicine, Inc. at the time the research was conducted. O.A.-W. has served as a consultant for H3B Biomedicine, Foundation Medicine Inc, Merck, Prelude Therapeutics, and Janssen, and is on the Scientific Advisory Board of Envisagenics Inc., AIChemy, Harmonic Discovery Inc., and Pfizer Boulder; O.A.-W. has received prior research funding from H3B Biomedicine, LOXO Oncology, and Nurix Therapeutics unrelated to the current manuscript. KA is a consultant for Syros Pharmaceuticals, is on the SAB of CAMP4 Therapeutics. KA and ZT received research funding from Novartis not related to this work.

## Data and materials availability

The raw and processed RNA-Seq data from cell lines used in this study are available on GEO (accession GSE 198518 – reviewer token available). Processed data for PDX samples used in this study are available on GEO (accession GSE 198518), raw data from PDX samples are not provided due to patient privacy concerns. Original code has been deposited at Zenodo and cited in the references for this paper. Any additional information required to reanalyze the data reported in this paper is available from the lead contact upon request.

## Notes

### Competing Interest Statement

The authors have declared no competing interest.

