## Supplemental data for "Splicing modulators impair DNA damage response and induce killing of cohesin-mutant MDS/AML"

### Supplementary Information

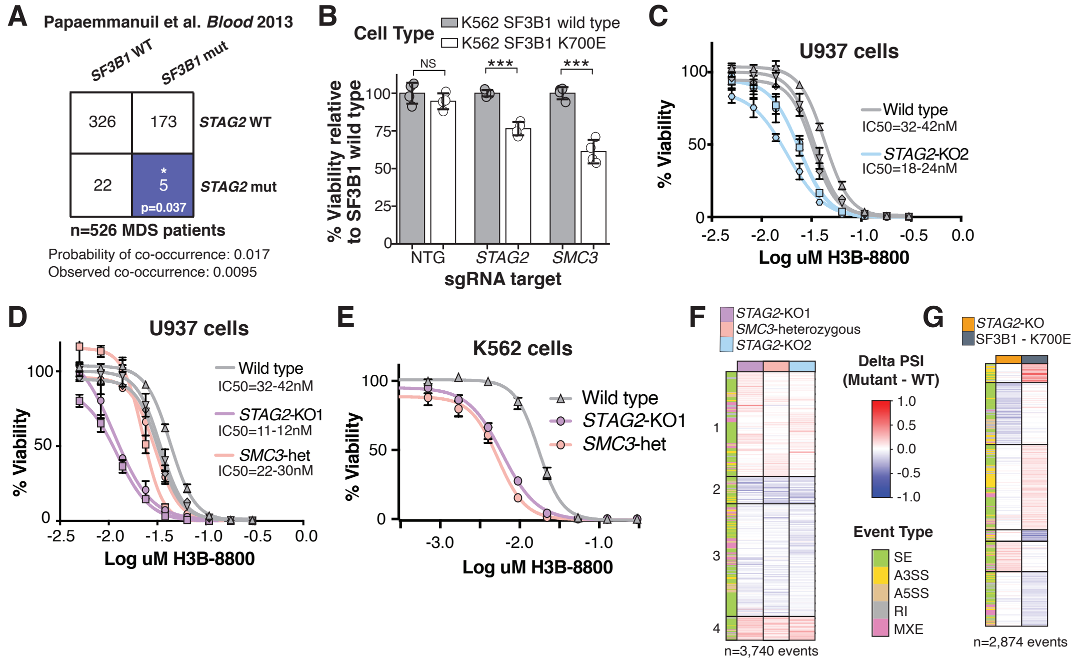

**Fig. S1. Cohesin-mutant cells are sensitive to SF3B1-targeting compounds.** (**A**) Co-occurrence of *SF3B1* and *STAG2* mutations in a cohort of MDS patients (*1*). Expected and observed probably of co-occurrence are listed. Blue color indicates a significant mutual exclusive relationship between *SF3B1* and *STAG2* mutations. *p<0.05 (Z-test). WT = wild type, mut = mutant. (**B**) Cell viability of wild type and SF3B1 K700E-mutant K562 cells following depletion of cohesin complex proteins with sgRNAs. Values are normalized to SF3B1 wild type cells for each knock-out or non-targeting guide RNA (NTG). Cell counts were taken 6 days after transduction with sgRNAs. Data shown are independent biological replicates and error bars represent +/- standard deviation. ***p<0.001 Student's t-test. (**C**,**D**) Drug dose-response curves of WT, *STAG2*-mutant, and *SMC3*-mutant U937 clones on day 8 of treatment with H3B-8800. Error bars represent SD of three technical replicates. (**E**) Drug dose response curves of one representative example of WT and *STAG2*-KO, and *SMC3*-het K562 clones on day 12 of treatment with H3B-8800. IC50 (WT) =20nM, IC50 (*STAG2*-KO) = 6nM, IC50 (*SMC3*-het) = 5nM. Error bars represent SD of measurements of three technical replicates. (**F**) k-means clustering of delta percent spliced in (PSI) scores for all events detected in any comparison of a cohesin-mutant genotype to wild type controls in U937 cells. Each row represents a splicing event that is significantly regulated in at least one condition, and the values plotted are delta PSI (mutant – wild type) scores averaged from independent single-cell clones in each condition. Color bar on the left indicates the type of splicing event that was called. (**G**) k-means clustering of delta PSI scores for all significant splicing changes (delta PSI > 5%, FDR < 0.05) detected in SF3B1 K700E-mutant (*2*) or *STAG2*-KO K562 cells (*3*). Heatmap values are as described in panel 1F.

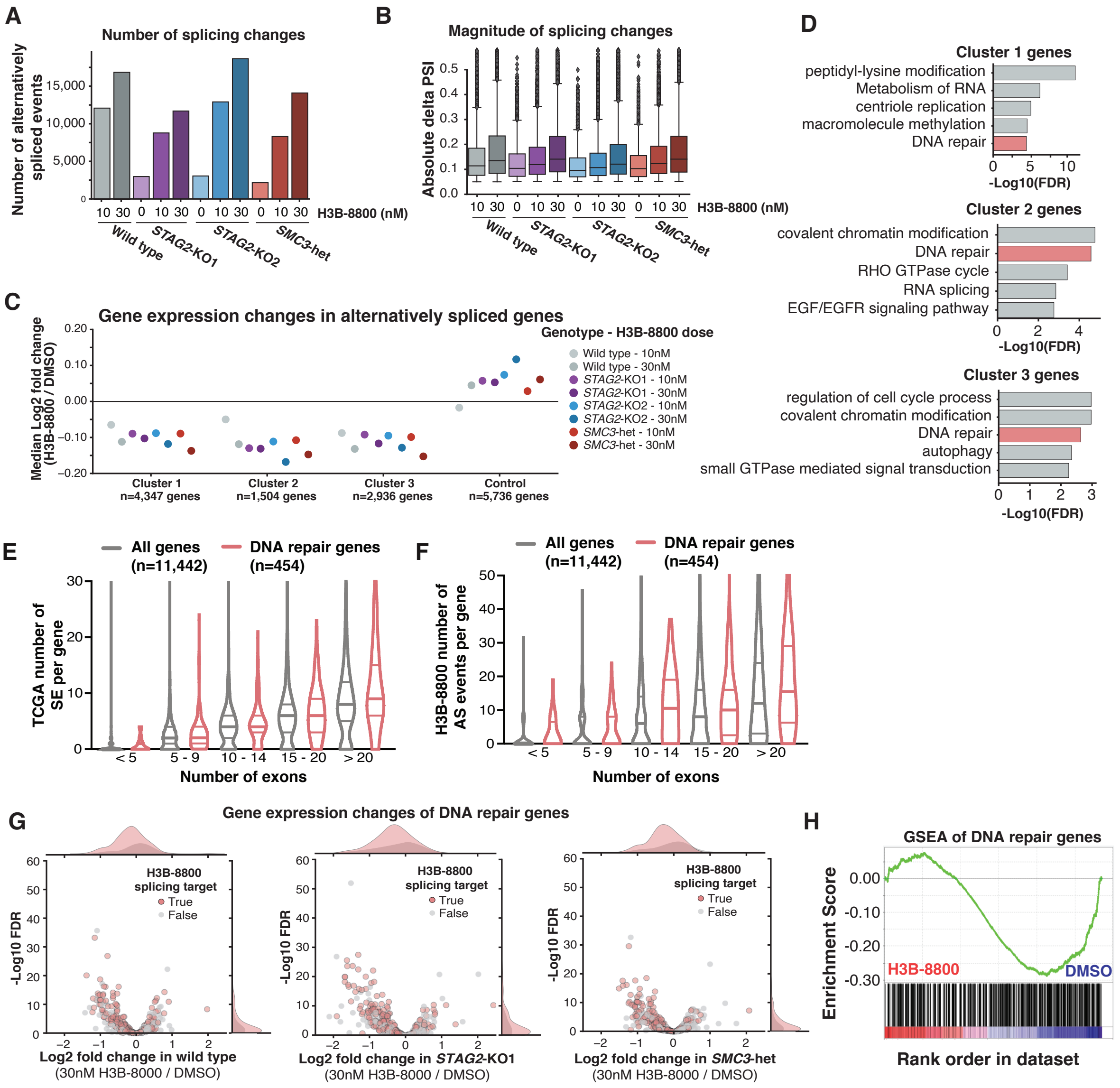


**Fig. S2. H3B-8800 treatment induces mis-splicing and downregulation of DNA damage repair genes.** (**A**) Total number of significant splicing changes (FDR < 0.05, delta PSI > 5%, 2 or 3 biological replicates included per condition) in each mutant genotype and drug treatment. Cohesin-mutant cell lines that were treated with DMSO (0nM H3B-8800) are compared to DMSO-treated wild type cells. For each cell line treated with H3B-8800, the number of events is calculated compared to the DMSO-treated condition of the same genotype. (**B**) Absolute value of delta PSI (percent spliced in) found in all significant H3B-8800–regulated splicing changes (FDR < 0.05, delta PSI > 5%, 2 or 3 biological replicates included per condition). Cohesin-mutant cell lines that were treated with DMSO (0nM H3B-8800) are compared to DMSO-treated wild type cells. For each cell line treated with H3B-8800, the number of events is calculated compared to the DMSO-treated condition of the same genotype. Boxes represent the median value and the interquartile range. Whiskers extend to the rest of the distribution except outlier points that are plotted as diamonds. (**C**) Median log2 fold change of genes in each cluster of H3B-8800–regulated splicing changes. A control set was generated from genes that are not alternatively spliced in H3B-8800–treated cells. Fold change in gene expression was calculated for each concentration of H3B-8800 relative to DMSO-treated cells of the same genotype (n=2 or 3 biological replicates per genotype). (**D**) Top 5 gene ontology terms ranked on significance among alternatively spliced genes in each cluster of H3B-8800–regulated splicing events (Figure 2B). Ontology enrichment was performed with Metascape (*4*). (**E**,**F**) Violin plots comparing the number of skipped exons in TCGA (E) or the number of alternatively spliced (AS) events detected following H3B-8800 treatment (F) per gene by the number of exons in each gene. Comparing DNA repair genes (n=454) to all expressed protein-coding genes (n=11,442). Number of genes in each exon group are as follows (DNA repair genes in brackets). <5 exons: 2595 (34), 5-9 exons: 3733 (112), 10-14 exons: 2347 (102), 15-20 exons: 1396 (90), and >20 exons: 1371 (116). Horizonal lines in violin plots depict the median and 1$^{st}$ and third quartiles. (**G**) Volcano plot of gene expression changes in DNA repair genes in wild type, *STAG2*-KO1, and *SMC3*-heterozygous cells treated with 30nM H3B-8800, relative to DMSO-treated controls. DNA repair genes that contain H3B-8800–regulated splicing changes are highlighted in red. (**H**) Gene set enrichment analysis (GSEA) of curated list of DNA repair proteins (*5*) comparing all H3B-8800–treated cell lines (n=20) against DMSO-treated controls (n=10). Normalized enrichment score = -1.1, q-value = 0.375.

A

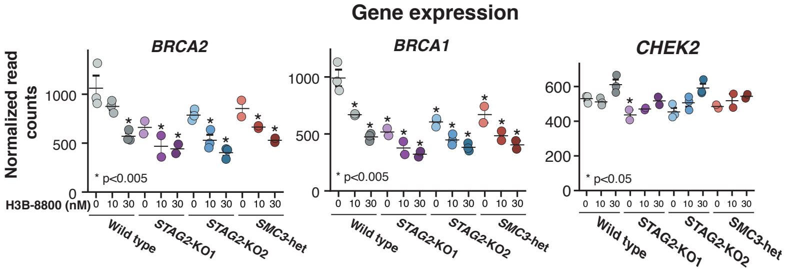


**Fig. S3. Mis-splicing of DNA repair genes alters protein function and results in accumulation of DNA damage.** (**A**) DESeq2 normalized read counts measuring gene expression of *BRCA2, BRCA1,* and *CHEK2*. Error bars represent mean +/- SEM of biological replicates (n=2 or 3 per genotype). *p<0.005 relative to DMSO-treated wild type cells (FDR-corrected Wald test).

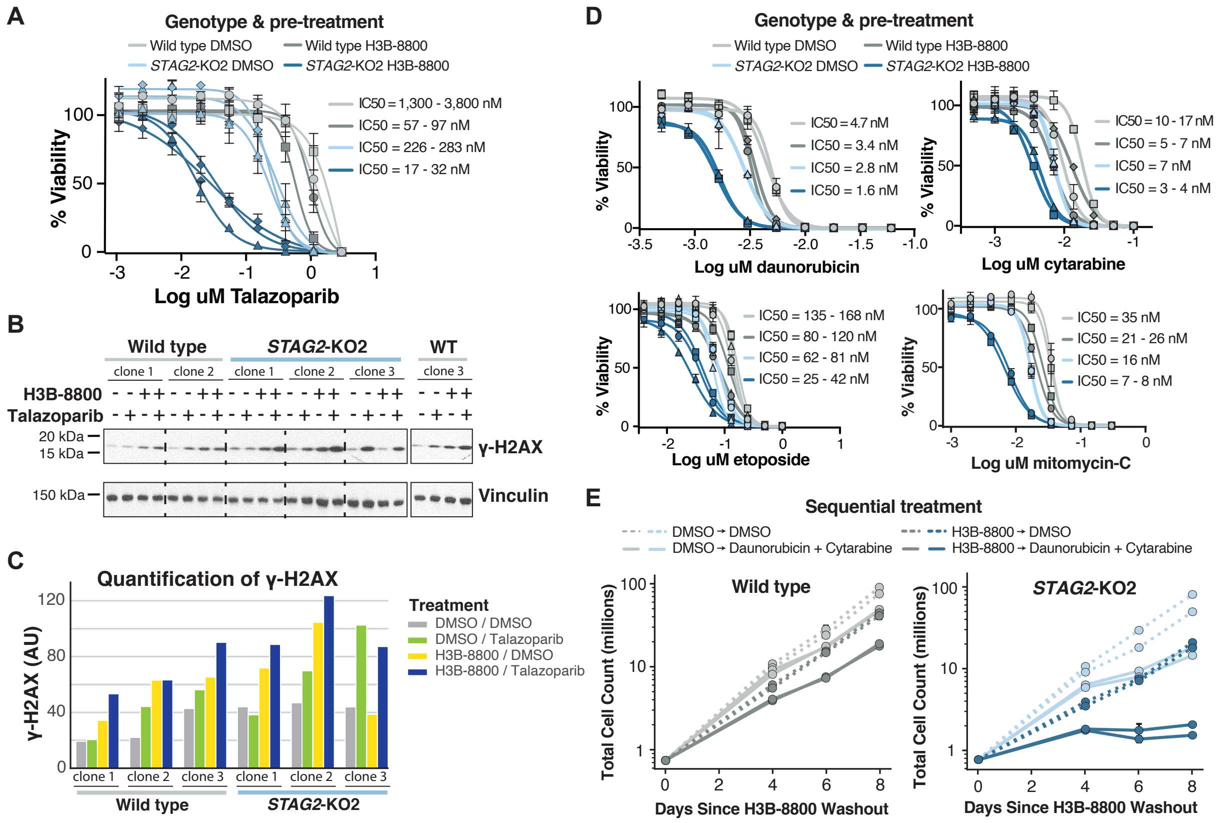


**Fig. S4. Splicing modulation sensitizes cohesin-mutant AML cell lines to killing by talazoparib and chemotherapy.** (**A**) Drug dose-response curves of wild type and *STAG2*-KO2 cells pre-treated for 3 days with 1.14 nM E-7107 or DMSO, followed by 8 days of treatment with talazoparib. Error bars represent SD of technical triplicate measurements for each biological replicate sample (n=2 per genotype and condition). (**B**) Western blot analysis of γH2AX in individual U937 single-cell clones after sequential treatment with H3B-8800 or DMSO and talazoparib or DMSO. Cells were treated with either H3B-8800 or DMSO for 3 days (top row), when the drug was washed out and replaced with either talazoparib or DMSO for 4 days (bottom row). Cells were collected for western blot on day 7. (**C**) Quantification of γH2AX western blot (panel B) summarizing independent single-cell clones of each genotype. (**D**) Drug dose-response curves of wild type and *STAG2*-KO2 cells pre-treated with 50nM H3B-8800 for 3 days followed by 8 days of treatment with daunorubicin, cytarabine, etoposide, or, mitomycin-C. Error bars represent SD of measurements in technical triplicates measurements for each biological replicate sample (n=2 per genotype and condition). (**E**) Growth curves depicting total number of wild type or *STAG2*-KO2 cells pre-treated with DMSO or H3B-8800 (50nM) into 8 days of DMSO or daunorubicin (5nM) + cytarabine (10nM) combination treatment. Error bars represent SD of technical duplicate measurements for each biological replicate sample (n=2 per genotype and condition).

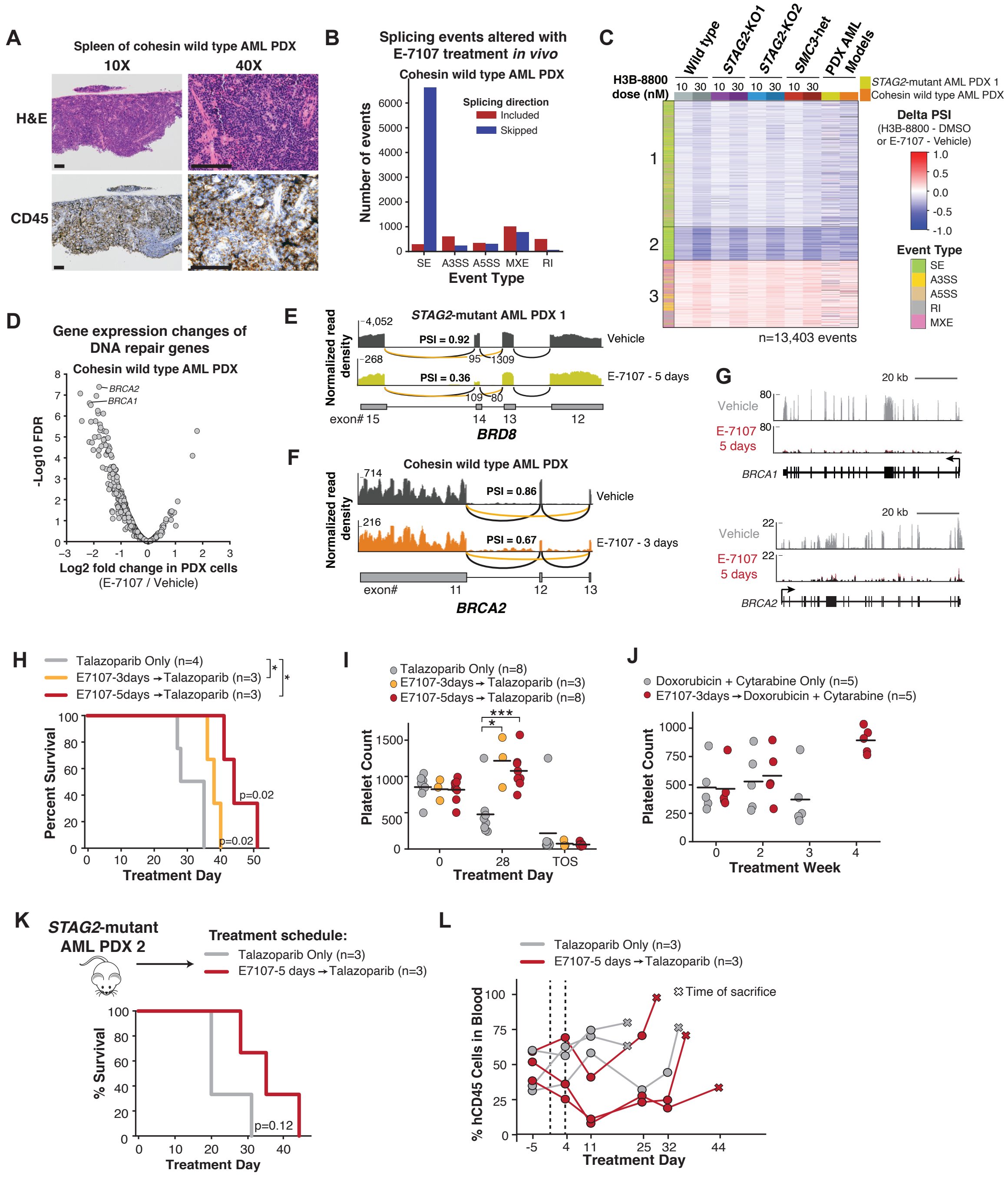

**Fig. S5. Low-dose splicing modulation combined with talazoparib targets PDX AML *in vivo.***

(**A**) Morphologic evaluation of spleen of cohesin wild type AML patient derived xenograft shows infiltration with human leukemia blasts. Images were stained using H&E and hCD45-targeting antibody and imaged at 10x and 40x (scale bar: 0.125mm) original magnification. (**B**) Total number and directionality of significant splicing alterations differentially called in cohesin wild-type human AML cells isolated from bone marrow of NSGS mice treated with E-7107 compared to vehicle for 3 days *in vivo.* Splicing events are categorized by event type and direction of regulation in E-7107 versus vehicle treated mice. (SE = skipped exon, A3SS = alternative 3' splice site, A5SS = alternative 5' splice site, MXE = mutually exclusive exon, RI = retained intron). n=2 mice per condition. (**C**) Heatmap of delta PSI scores for H3B-8800–regulated exons called from U937 cells (Figure 3A) including two human AML PDX models that were sequenced. Delta PSI scores from PDX models compare E-7107 to vehicle treated control mice for 3 days (cohesin wild type AML) or 5 days (*STAG2*-mutant AML). n=3 mice per condition for *STAG2*-mutant AML and n=2 mice for cohesin wild type AML. Color bar on the left indicates the type of splicing event that was called. (**D**) Volcano plot depicting differential gene expression of DNA repair genes in cohesin wild type AML PDX cells isolated from the bone marrow of NSGS mice treated with E-7107 versus vehicle for 3 days *in vivo*. n=2 mice per condition. (**E**) RNA-Seq normalized read density and splice junction track of exon skipping in *BRD8* exon 14 from one representative replicate of vehicle treated *STAG2*-mutant PDX AML cells and the combined coverage of 3 replicates of E-7107 treated cells. Average PSI scores of exon14 are shown. Average number of reads supporting exon skipping (orange line) and exon inclusion (black line) are reported. PSI (percent spliced in). (**F**) RNA-Seq normalized read density and splice junction track of exon skipping in *BRCA2* exon 14 from one representative replicate of vehicle and E-7107 treated cohesin wild type AML PDX model. Average PSI scores of exon12 are shown. Constitutive junctions are shown with black lines, exon skipping junctions are shown in orange. PSI (percent spliced in). (**G**) Browser tracks of RNA-Seq coverage on *BRCA1* and *BRCA2* genes from *STAG2*-mutant human AML1 PDX cells isolated from the bone marrow of NSGS mice treated with E-7107 or vehicle for 5 days *in vivo*. 3 mice per condition are merged for visualization. (**H**) Survival analysis from first *in vivo* experiment with E-7107 and talazoparib for *STAG2*-mutant AML1 PDX model (other mutations include *BCOR/RUNX1/U2AF1/DNMT3A*). Treatment of mice assigned to three treatment arms was initiated 3 weeks after bone marrow transplantation: talazoparib only (n=4), E-7107 for 3 days followed by talazoparib (n=3), or E-7107 for 5 days followed by talazoparib (n=3). P-values shown are compared to the talazoparib only arm (log-rank test). (**I**) Platelet count from peripheral blood of *STAG2*-Mutant AML1 PDX mice before treatment (T0), 4 weeks after start of treatment (4wk), and at the time of sacrifice (TOS). Students t-test comparing each treatment group to talazoparib-only group for each timepoint. (**J**) Platelet count from peripheral blood of *STAG2*-Mutant AML1 PDX mice before treatment (Week 0) and at each week while on treatment. (**K**) Survival analysis of *STAG2*-mutant AML2 PDX model (other mutations include *RUNX1, ASXL1, NRAS*) treated with either talazoparib alone, or E-7107 for 5 days followed by talazoparib until the time of sacrifice. P-value is calculated with a log-rank test. (**L**) Percent of hCD45+ cells in the peripheral blood throughout treatment until time of sacrifice. Each line represents an individual mouse used in the experiment. Dashed lines indicate the 5-day treatment window with E-7107 or vehicle control. %hCD45 cells at the time of sacrifice are indicated with an X.

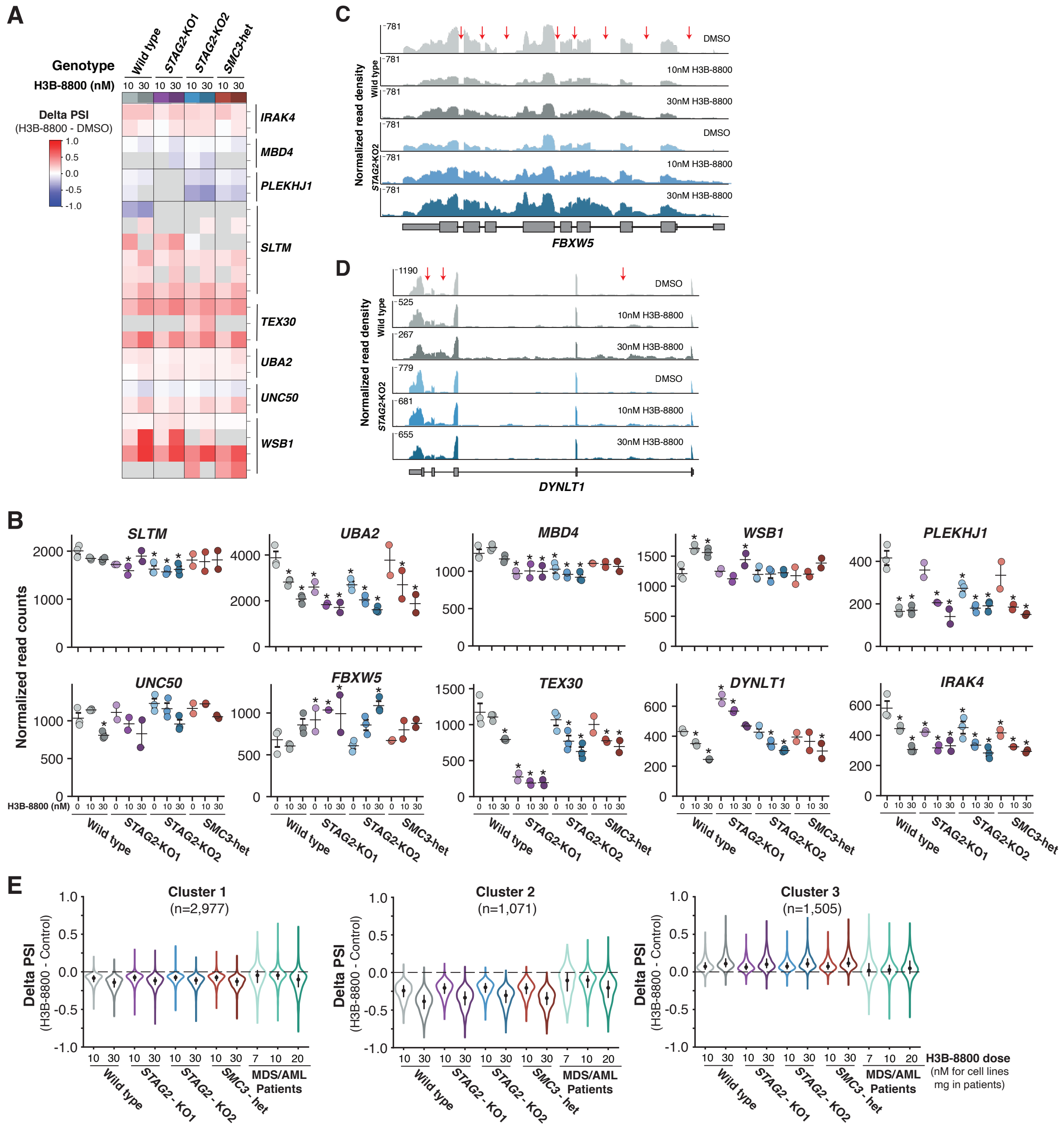

**Fig. S6. Splicing changes and downregulation of DNA repair genes are conserved in MDS/AML patients.** (**A**) Delta PSI of all H3B-8800–regulated splicing events detected in genes that were used as biomarkers of on-target activity in the Phase 1 clinical trial (*6*). Delta PSI is the average difference of 2 or 3 biological replicates per drug concentration relative to DMSO-treated controls of the same genotype. Events that were detected in at least one comparison are included in the heatmap. Grey boxes indicate the event was not detected in the represented sample. (**B**) DESeq2 normalized read counts of the genes used as biomarkers of on-target activity in the Phase 1 clinical trial of H3B-8800 that are expressed in U937 cells. Error bars represent mean ± SEM of biological replicates (n=2 or 3 per genotype). *FDR $< 0.05$ compared to DMSO-treated wild type cells (FDR-corrected Wald test). (**C**) RNA-Seq normalized read density from one representative example gene (*FBXW5*) in H3B-8800–treated wild type and *STAG2*-KO2 cells. Red arrows indicate regions with a dose-dependent effect on intron retention. (**D**) RNA-Seq normalized read density from one representative example gene (*DYNLT1*) in H3B-8800–treated wild type and *STAG2*-KO2 cells. Red arrows indicate regions with a dose-dependent effect on intron retention. (**E**) Violin plots depicting the Delta PSI scores for splicing events in each cluster from Figure 5C. Dot represents the median, and bars extend from the first to third quartile range.

**Table S1**. **Gene Ontology of H3B-8800 Target Genes.** Metascape summary for enrichment of gene ontology terms among genes that are mis-spliced in each cluster (Figure 2B) in cells treated with H3B-8800. The top 20 enriched categories are reported for each cluster of genes, ranked on significance value. In cases where more than 500 genes were in a cluster, the top 500 ranked on magnitude of deltaPSI were used to calculate enrichment of gene ontology categories.

| Patient | Sex | Disease Type | Karyotype & Mutation | H3B-8800 drug dose |
|---|---|---|---|---|
| #1 | Male | MDS EB2 | 46XY, IDH1 132C, SRSF2 P95H | 7mg po daily |
| #2 | Male | AML MRC transformed from RAEB-2 | 47 XY, +19[7], SRSF2 P95H, DNMT3A R882H, BCOR R1661*, SETD1B E1146delinsV, NRAS G12A | 10mg po daily |
| #3 | Male | AML transformed from CMML-2 | 46 XY, +8 [20], ASXL1 L823*, RUNX1 W108fs*33, SRSF2 P95L | 20mg po daily |

**Table S2**. **Patient Sample Information.** Information on disease type and mutation status of patients for whom sequencing was performed on peripheral blood samples pre- and post- H3B-8800 treatment.
